# The histone variant H2A.Z and its chaperone SRCAP are required for RNA polymerase II to access HBV chromatin

**DOI:** 10.1101/2025.11.10.687638

**Authors:** Basile Jay, Océane Lopez, Laetitia Gérossier, Nazim Sarica, Gabrielle Pozo, Sabine Gerbal-Chaloin, Martine Daujat-Chavanieu, Olivier Hantz, Christine Neuveut, Joao Diogo Dias

## Abstract

**Background:** Chronic hepatitis B virus (HBV) infection remains a major global health burden due to the persistence of covalently closed circular DNA (cccDNA), a stable episomal viral reservoir resistant to current antiviral therapies. The transcriptional activity of cccDNA depends on its chromatin organization, yet the contribution of histone variants to this regulation remains poorly understood.

**Methods:** Using mass spectrometry–based proteomics on native cccDNA purified from infected primary human hepatocytes, we identified the histone variant H2A.Z and its chaperone SRCAP as cccDNA-associated proteins. Functional analyses combining shRNA-mediated silencing, chromatin immunoprecipitation (ChIP), and ATAC-seq were performed in HBV-infected HepG2-NTCP cells and primary human hepatocytes to characterize their roles in cccDNA formation and transcription.

**Results:** Depletion of H2A.Z.1, H2A.Z.2, or SRCAP reduced both HBV RNA levels and cccDNA formation. H2A.Z recruitment to cccDNA correlated with the establishment of active chromatin marks (H3K4me3) and enhanced RNA polymerase II loading, promoting an open and transcriptionally active chromatin state. In addition, we identified BRD2, an H2A.Z-associated transcriptional co-activator, as a positive regulator of HBV transcription. shRNA mediated depletion and pharmacological degradation of BRD2 using the PROTAC ARV-771 reduced HBV RNA levels, supporting its potential as an antiviral target.

**Conclusion:** Our findings uncover a critical role of the H2A.Z variant and SRCAP complex in cccDNA formation and transcriptional activation. By facilitating chromatin accessibility and RNA polymerase II recruitment, H2A.Z establishes an epigenetic environment favorable to HBV persistence. Targeting H2A.Z-associated co-activators such as BRD2 may represent a promising strategy to silence cccDNA transcription and achieve functional HBV cure.

## INTRODUCTION

Hepatitis B virus (HBV) is a widespread pathogen and one of the most important environmental risk factors in human cancer epidemiology^1^. Despite the existence of an effective vaccine, the number of chronic HBV-carriers remains elevated with 350 million individuals worldwide at increased risk of developing severe liver diseases such as liver fibrosis, cirrhosis and hepatocellular carcinoma (HCC). Current treatments for chronic hepatitis B, including nucleos(t)ide analogues and interferon-α, control viral replication and improve liver function; however, because they cannot eliminate the episomal nuclear viral DNA known as covalently closed circular DNA (cccDNA), they fail to achieve complete viral clearance. Consequently, a major area of research focuses on understanding cccDNA biology to develop new therapeutic strategies capable of neutralizing this viral reservoir.

HBV is a 3.2 kb DNA virus that replicates its genome in the cytoplasm through reverse transcription of the encapsidated pregenomic RNA (pgRNA) into a partially double-stranded circular DNA (RC-DNA)^2^. Upon internalization of the viral particles, the capsid delivers RC-DNA to the nucleus, where it is converted into cccDNA. This episomal molecule represents a key intermediate in the HBV life cycle. First, it serves as a template for the transcription of all viral RNAs including the pgRNA which in turn replenishes the pool of cccDNA. Second, cccDNA is a long-lived intermediate that maintains viral genetic information, acting as a viral reservoir in infected individuals.

The establishment of a transcriptionally active cccDNA molecule is a multistep process requiring host-cell DNA synthesis and repair factors to convert RC-DNA into covalently closed viral DNA, followed by engagement of the cellular transcriptional machinery^3^. Within the nucleus, cccDNA is organized similarly to cellular DNA, associated with the four core histones (H2A, H2B, H3, and H4) and the linker histone H1^4,5^, forming a chromatin-like structure containing nucleosomes. The precise timing of histone deposition on viral DNA, whether occurring on RC-DNA or only after cccDNA formation, remains unclear, but a recent study suggest that cccDNA chromatinization occurs concomitantly with cccDNA formation^6^. Finally, multiple studies have confirmed the presence of histones on cccDNA both in infection models and in vivo, and demonstrated their role in HBV transcriptional regulation^7–18^. Recently Prescott and collaborators using in vitro reconstituted cccDNA highlighted the importance of nucleosome organization for proper HBV transcription^11^.

In addition to canonical histones expressed exclusively during the S-phase of the cell cycle, different histone variants, whose expression is cell cycle independent, can replace them, diversifying chromatin structure and contributing to more specialized functions. Recent studies identified histone variant H3.3 as a positive regulator of both cccDNA formation and transcription^6,19^. Using silencing approaches they further demonstrated that the histone regulator A (HIRA) complex is required for its deposition^6^. Among histone variants, the H2A family comprises several members including macroH2A, H2AX and H2A.Z. H2A.Z is a replication independent histone with two isoforms in mammals: H2A.Z.1 and H2A.Z.2, the latter comprising two splice variants in primates: H2A.Z.2.1 and H2A.Z.2.2. H2A.Z has been implicated in both transcriptional activation and repression, depending on the chromatin context and associated post-translational modifications. H2A.Z typically accumulates upstream and downstream of transcription start sites (TSSs) being enriched at the +1 nucleosome within gene bodies and at enhancers of active or transcriptionally poised genes^20–24^. Recent studies suggest that H2A.Z incorporation enhances nucleosome dynamics and DNA unwrapping, thereby increasing DNA accessibility to the transcriptional machinery and chromatin modifying factors^25–27^. Beyond transcription, H2A.Z has also been associated with DNA repair, replication, chromosome segregation, and cell-cycle progression^21,25,28–30^. Although highly similar in sequence, recent evidence indicates that the two H2A.Z isoforms are not entirely functionally redundant and display distinct chromatin distributions and roles across different tissues^22,29,31^. Interestingly, H2A.Z activity depends on its nucleosomal context, particularly on the presence of other histone variants and specific post-translational modifications. Notably, nucleosomes containing both H3.3 and H2A.Z variants mark nucleosome free regions at active promoters and enhancers, facilitating the access to transcription factors^32,33^. A study conducted in mouse embryonic stem cells (mESCs) also highlighted cooperation between the HIRA complex and SRCAP in depositing H2A.Z at poised genes^34^.

Although recent studies have clearly established the importance of proper chromatin organization and histone post-translational modifications in regulating HBV transcription, much less is known about the contribution of histone variants to this process. In this study, using a method allowing the purification of native cccDNA from infected primary human hepatocytes, followed by mass spectrometry analysis of associated proteins, we identified the histone variant H2A.Z and its chaperone SRCAP. Using loss-of-function approaches in different cellular HBV infection models, we demonstrated that H2A.Z.1, H2A.Z.2, and SRCAP are required for both HBV cccDNA formation and transcription. We further showed that H2A.Z deposition is correlated with the establishment of an open chromatin state, thereby facilitating the recruitment of RNA polymerase II. Finally, we demonstrated that Brd2, an H2A.Z-associated transcriptional activator, is recruited to cccDNA, where it contributes to HBV transcriptional activation and may represent a promising therapeutic target.

## RESULTS

### Identification of H2A.Z histone variant and SRCAP as cccDNA associated proteins

To identify cellular proteins associated with cccDNA, we purified cccDNA from infected primary human hepatocytes using an iodixanol gradient, followed by mass spectrometry analysis. We identified several proteins previously reported to associate with cccDNA, including HNF4, RNA polymerase II, TFIIH, PRMT5, and the core histones H3, H4, H2A, and H2B. In addition, we identified new proteins, notably the histone variant H2A.Z and the SNF2-related CREBBP activator protein (SRCAP), a component of the SRCAP chromatin-remodeling complex that functions as an H2A.Z chaperone (Supp. Table 1). The presence of these functionally linked proteins on cccDNA in infected cells suggests that this histone variant may play an important role in the HBV replication cycle. The recruitment of H2A.Z to cccDNA was further confirmed by cccDNA ChIP-qPCR in wild type HBV (HBVwt) infected HepG2-NTCP-sec and Primary Human Hepatocytes (PHH) cells using an antibody that recognizes both H2A.Z variants (Fig. 1B, Supp. Fig. 1A). H2A.Z deposition has been associated with both transcriptional activation and repression, while hyperacetylation of H2A.Z is typically linked to gene activation^35–38^. We next demonstrated that cccDNA is enriched in acetylated H2A.Z, suggesting that H2A.Z is involved in HBV transcriptional activation (Fig. 1C). Furthermore, both H2A.Z and acetylated H2A.Z are present in similar proportions across the viral promoters in HepG2 (Supp. Fig. 1B, C), suggesting a global effect on viral transcription.

**Figure 1:**
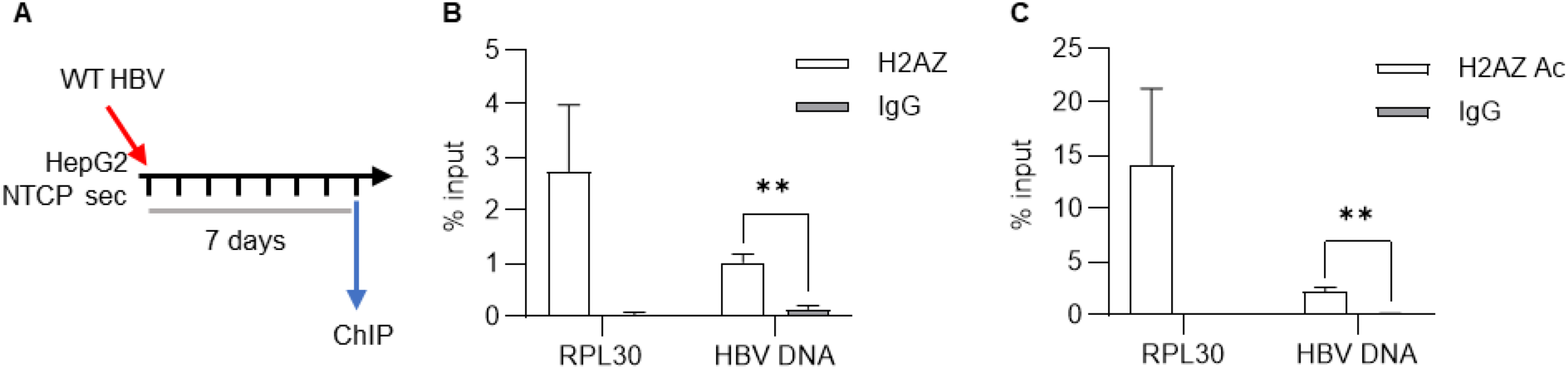
H2AZ is recruited to HBV cccDNA chromatin. (A) HepG2 NTCP sec cells were infected with WT HBV at an MOI of 200 geq/cell for 7 days before chromatin extraction. (B-C) ChIP assays were performed on MNAse fragmented chromatin using anti-H2AZ antibody, anti-acetylated H2AZ antibody or nonspecific rabbit IgG as indicated and followed by qPCR analysis with primer specific for the enhancer 1 region or cellular region (RPL30). Mean ± SD of at least 3 experiments is shown. *P* values were determined by unpaired Student’s *t* test with Welch correction (**, P < 0.01).

### H2A.Z1 and H2A.Z2 histone variants and SRCAP are involved in cccDNA establishment and required for cccDNA transcription

To further investigate the role of H2A.Z on HBV RNA expression we silenced the two isoforms using shRNAs specifically targeting either H2A.Z.1 or H2A.Z.2. HepG2-NTCP-sec cells were first transduced with lentiviruses expressing the different shRNAs individually or in combination, and infected with HBV 48 hours later (Fig. 2A). Downregulation of H2A.Z.1 or H2A.Z.2 leads to a decrease in pgRNA and total HBV RNA levels (Fig. 2B, Supp. Fig. 2 A, B). Analysis of published transcriptomic data from HepG2 cells silenced or not for H2A.Z, showed that H2A.Z silencing did not significantly affect the expression of known HBV transcriptional regulators, except for JUN, whose expression was increased in H2A.Z-silenced cells (Supp Fig. 3 A). Similarly, we assessed the role of the chaperone SRCAP complex using shRNA-mediated knockdown and observed that SRCAP depletion correlates with a strong decrease in HBV RNA levels (Fig. 2C, Supp. Fig. 2C) without significantly affecting the expression of HBV-regulating transcription factors such as HNF4, HNF1, or CREB (Supp. Fig. 3B). The role of the SRCAP complex in HBV transcription was further confirmed by silencing VPS72, a key subunit of the complex, that equally resulted in a reduction of HBV RNA levels (Fig. 2D, Supp. Fig. 2D). Altogether, our results suggest that SRCAP complex is required for HBV transcription and may, at least in part, act through the deposition of H2A.Z on cccDNA as VPS72 is the subunit that directly interacts with and mediates the deposition of H2A.Z.

**Figure 2:**
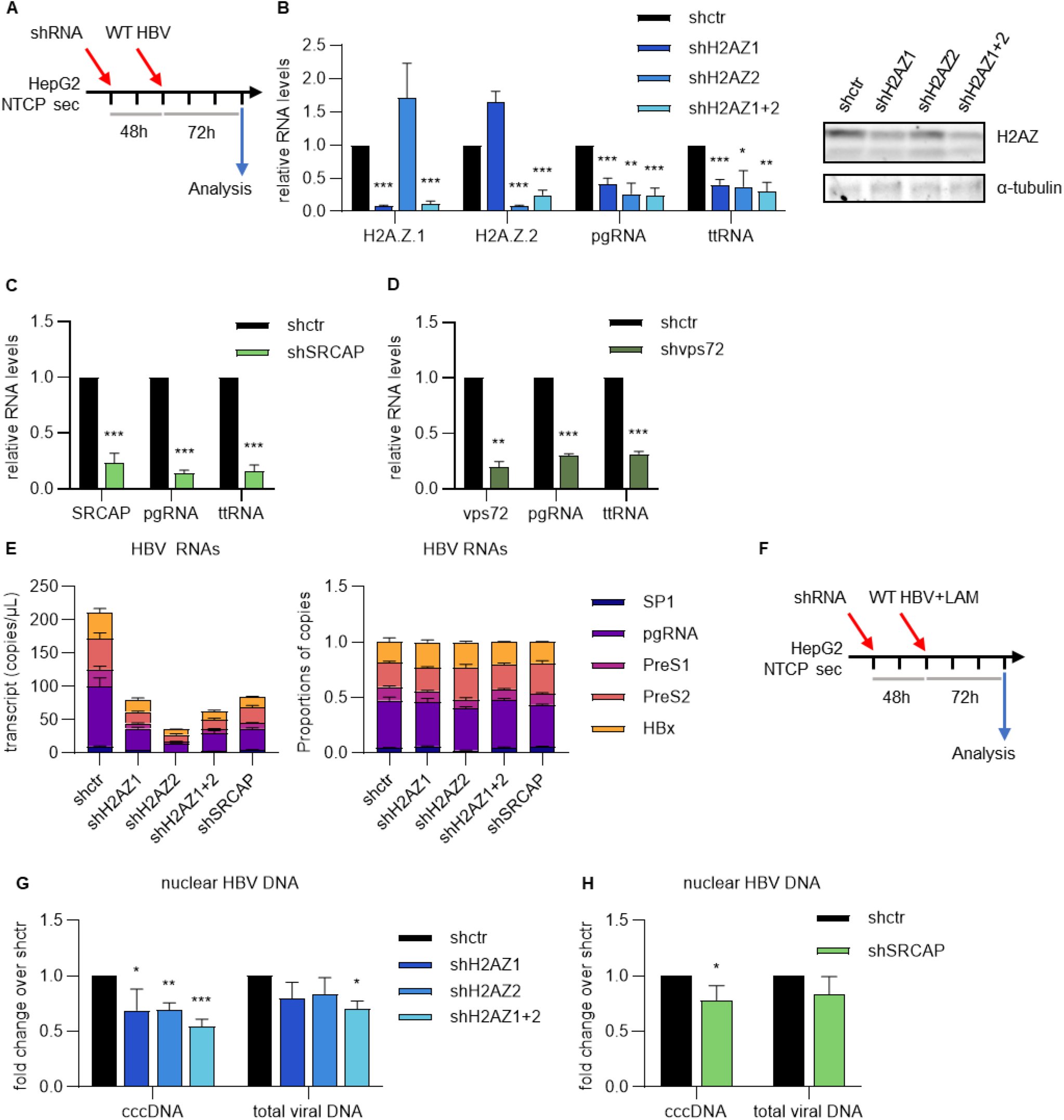
H2AZ and SRCAP are required for both cccDNA establishment and HBV RNAs expression in HepG2-NTCP cells. (A) HepG2 NTCP sec cells were transduced with shRNAs targeting H2AZ.1 or H2AZ.2 individually or in combination, SRCAP, or targeting VPS72 or shRNA control (shctr) for 48 hours before being infected with WT HBV at an MOI of 100 geq/cell. (B) to (F) Total RNAs and proteins were extracted at 72 hours post infection for analysis. Messenger RNA (mRNA) levels of H2AZ.1 or H2AZ.2 (B), SRCAP (C) or VPS72 (D) as well as HBV RNA levels: pgRNA and total HBV RNA (ttRNA) were analyzed by RT-qPCR normalized by the level of RHOT2 mRNA. Expression level in cells transduced with shRNA control was set to 1. Data represents mean ± standard deviation for at least 3 independent experiments, p value was calculated with an unpaired Student’s *t* test with Welch correction (*, P<0.05; **, P<0.01; ***, P<0.001). (B) H2AZ protein expression in HepG2 NTCP cells transduced by the indicated lentiviral constructs were analyzed by Western blot. Tubulin was used as loading control. (E) RNA levels of HBx RNA, PreS2/S RNA, PreS1 RNA, PreC/pgRNA and SP1 RNA were quantified by ddOTs using the same amount of total RNA. Data represents mean ± standard deviation for at least 3 independent experiments. Absolute quantification (left panel) or relative proportion of each transcript (right panel) are shown. (F) HepG2 NTCP sec cells were transduced with shRNAs targeting H2AZ.1 or H2AZ.2 individually or in combination, or targeting SRCAP or shRNA control (shctr) for 48 hours before being infected with WT HBV at an MOI of 100 geq/cell in the presence of lamivudine at 25µM. Nuclear DNA was extracted at 72 hours post infection. HBV cccDNA and total viral DNA were quantified by qPCR in cells silenced for H2AZ1 or H2AZ2 or both (G) or for SRCAP (H). Viral DNA quantification was normalized to cell numbers by qPCR amplification of the promoter of cyclin A2 (CCNA2). Viral DNA level in cells transduced with shRNA control was set to 1. Data represents mean ± standard deviation for at least 3 independent experiments, p value was calculated with an unpaired Student’s *t* test with Welch correction (*, P<0.05; **, P<0.01; ***, P<0.001).

HBV gene expression is controlled by four different promoters and two enhancers. We therefore assessed whether H2A.Z variants differentially affect HBV promoter activity using the recently described ddOTs approach^39^. Our results showed that all HBV promoters are similarly affected by H2A.Z or SRCAP silencing (Fig. 2E).

To rule out the possibility that viral gene silencing resulted from SMC5/6-induced transcriptional repression, we also examined the effect of H2A.Z knockdown in cells silenced for SMC6 expression (Supp. Fig. 4). Since in the absence of SMC5/6 complex the decrease in viral RNA levels is still observed we confirm that H2A.Z deposition regulates transcription from all the HBV promoters.

As H2A.Z and SRCAP were silenced prior to infection in our experimental setting, the observed decrease in HBV RNA levels could result not only from transcriptional modulation but also from the deregulation of early steps of infection. In particular, H2A.Z histone variants are known to be involved in DNA repair; therefore, their silencing could affect cccDNA levels, whose formation depends on the activity of cellular proteins and DNA repair pathways. We found that the silencing of H2A.Z variants or SRCAP leads to a decrease in cccDNA levels, whereas the total amount of viral nuclear DNA that comprise protein-free rcDNA and cccDNA, remained unchanged (Fig. 2G and H). These results suggest that H2A.Z variants are important for the efficient conversion of rcDNA into cccDNA.

Of note, the decrease of cccDNA level is however not sufficient to explain the decrease of viral RNAs (Supp. Fig. 5). All together our results show that H2A.Z variants and their chaperone SRCAP are involved in both HBV cccDNA establishment and HBV transcriptional regulation. Finally, we assessed the role of H2A.Z and SRCAP in HBV RNA expression in primary human hepatocytes. We first infected PHH with HBVwt and then silenced H2A.Z.1, H2A.Z.2, or SRCAP 48 hours after infection (Fig. 3A) since we observed a reduction in infection efficiency the longer it was performed after plating the PHH. Knockdown of SRCAP or H2A.Z.2 leads to a decrease in HBV RNA levels (Fig. 3B and C) without significantly affecting the expression of known HBV transcriptional regulators (Supp. Fig. 3C). Interestingly, H2A.Z.1 silencing does not affect HBV RNA expression in PHH. However, because silencing is performed in non-dividing cells after infection, when HBV cccDNA is already established, wrapped with histones, we cannot, completely rule out that H2A.Z already loaded onto cccDNA could not be fully removed. As expected, because HBV cccDNA is already established when silencing is performed, we did not observe a variation of HBV DNA level (Supp. Fig. 6A and B).

**Figure 3:**
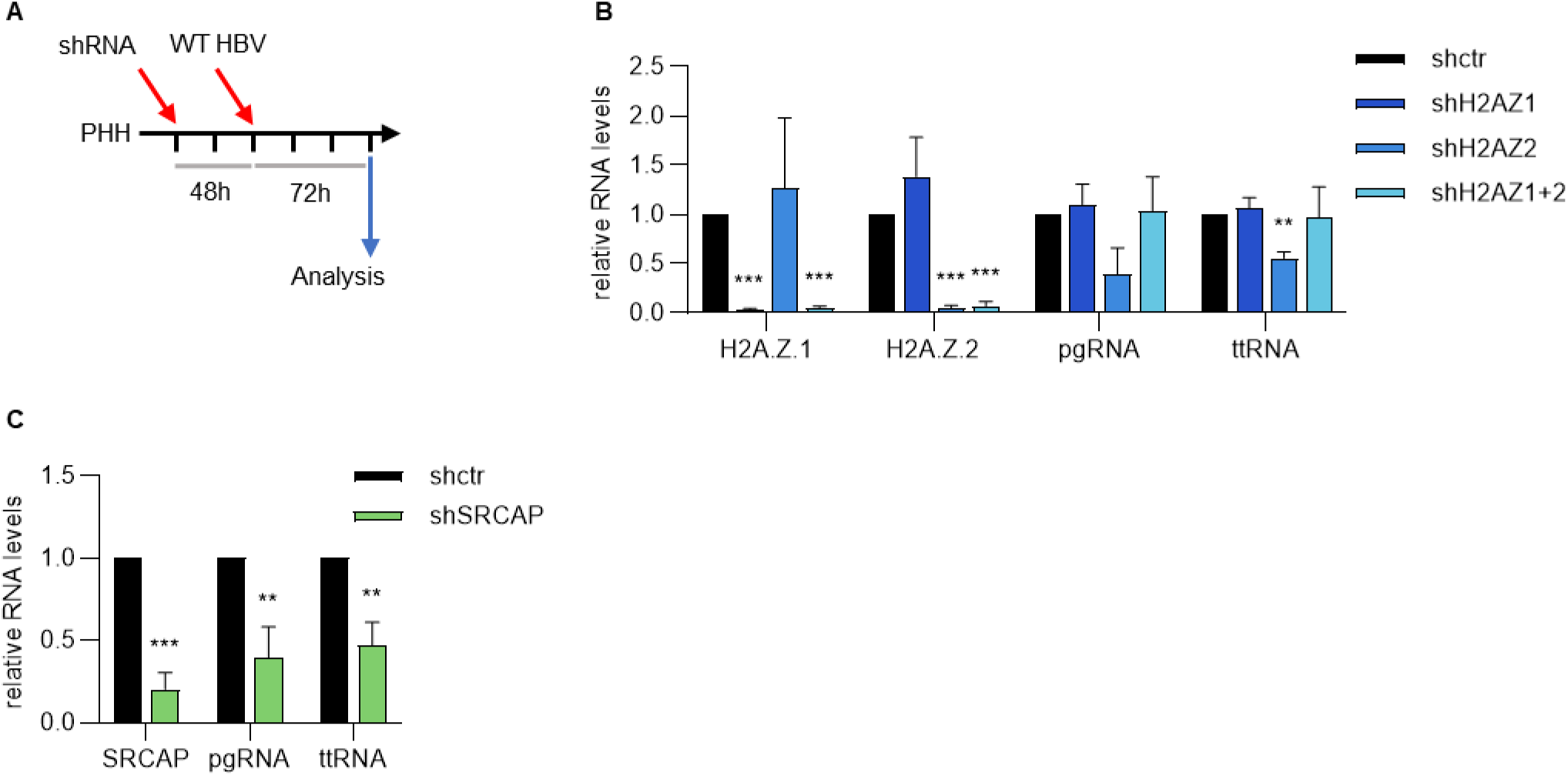
H2AZ and SRCAP are involved in HBV RNA transcription in primary human hepatocytes. (A) Primary human hepatocytes were infected with HBV at MOI of 100 geq/cells for 48 hours before being transduced with shRNAs targeting H2AZ.1 or H2AZ.2 individually or in combination or targeting SRCAP. RNA was extracted 5 days post infection. (B) and (C) mRNA levels of H2AZ.1 or H2AZ.2 (B), SRCAP (C) as well as HBV pgRNA and total HBV RNA (ttRNA) were evaluated by RTqPCR and normalized by the level of GAPDH mRNA. Expression levels in cells transduced with shRNA control was set to 1. Data represents mean ± standard deviation for at least 3 independent experiments, p value was calculated with unpaired Student’s *t* test with Welch correction (**, P<0.01; ***, P<0.001).

### H2A.Z and SRCAP regulate DNA accessibility and facilitate RNA Pol II loading to cccDNA

Having shown that silencing of H2A.Z or SRCAP leads to reduced HBV transcription, we next investigated the mechanism underlying H2A.Z-mediated transcriptional activation. H2A.Z incorporation has been shown to promote an open chromatin state, thereby increasing DNA accessibility to transcription factors and RNA polymerase II. We therefore first analyzed whether H2A.Z deposition modulates chromatin state and RNA polymerase II (RNAPII) recruitment to cccDNA. We first confirmed that H2A.Z is recruited to cccDNA and that SRCAP mediates its deposition, using ChIP-qPCR and MNAse digested chromatin from HepG2-NTCP-sec cells transduced with lentiviruses expressing shRNAs targeting H2A.Z.1, H2A.Z.2 or SRCAP and infected by HBV (Fig. 4A and B). We then analyzed histone post-translational modifications in cells silenced for H2A.Z or SRCAP. While we did not observe a strong decrease in H3K4me3 levels on viral chromatin, we nevertheless noted a downward trend in this mark. Silencing of H2A.Z.1, H2A.Z.2 or SRCAP was also not associated with an increase in repressive histone mark H3K9me3 (Fig. 4C and 4D). The GAPDH promoter was used as a control, since H2A.Z silencing does not affect GAPDH transcription (Fig. 4E, Supp. Fig. 8).

**Figure 4:**
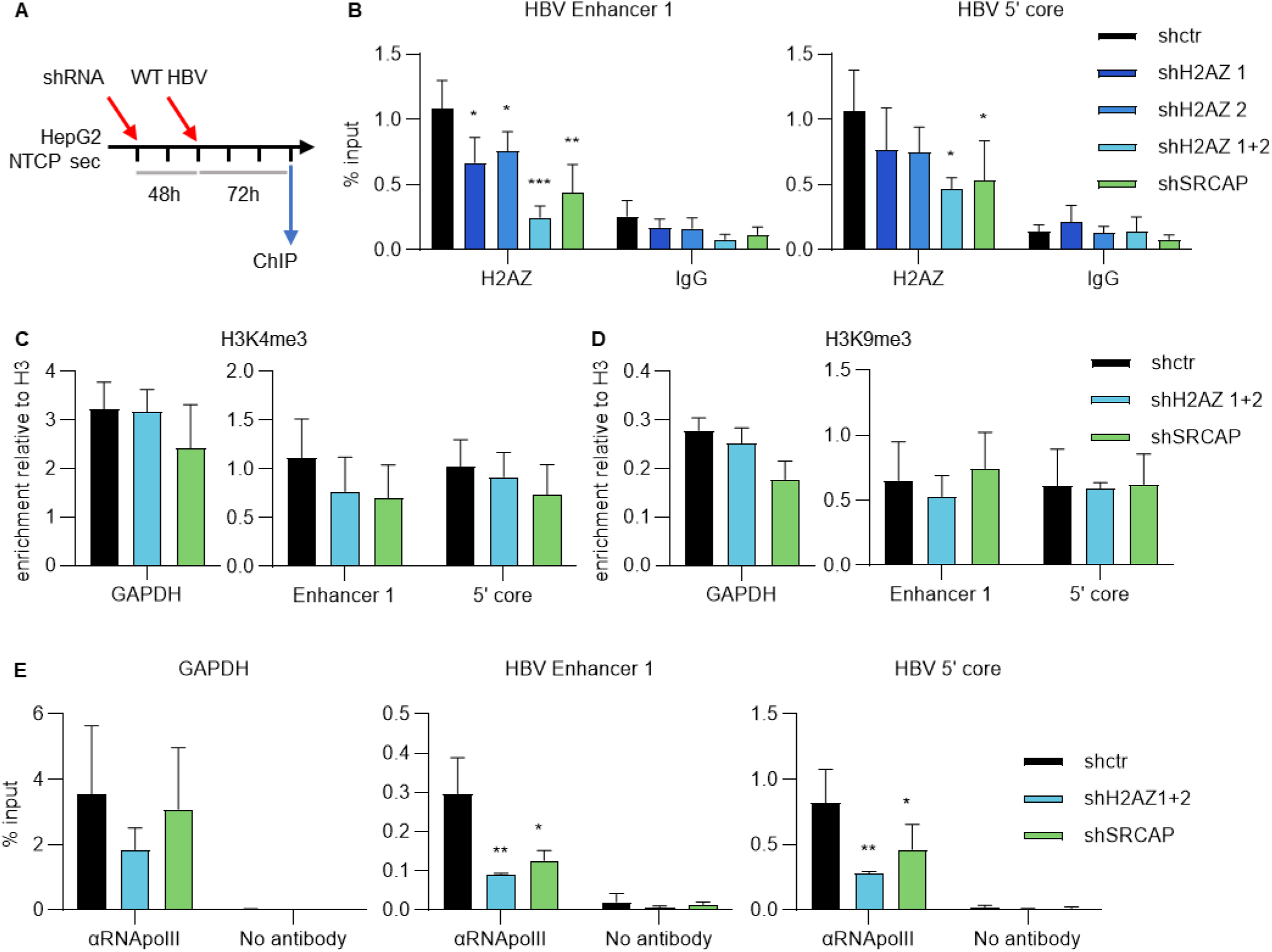
Knocking down H2AZ or its chaperone SRCAP impacts the levels of H2AZ and RNAPII on the viral chromatin. (A) HepG2 NTCP sec cells were transduced with shRNAs targeting H2AZ.1 or H2AZ.2 or both or targeting SRCAP for 48 hours before being infected with WT HBV at an MOI of 200 geq/cell. The cells were crosslinked 72 hours post infection before chromatin extraction. (B) to (D) MNAse fragmented chromatin was immunoprecipitated with antibody targeting H2AZ (B), H3 (C-D), H3K4me3 (C), H3K9me3 (D), and nonspecific IgG (B) and analyzed by qPCR using primers specific for HBV enhancer 1 region or for the 5’ core promoter region or the GAPDH promoter region. The levels of enrichment for H3K4me3 and H3K9me3 were normalized by the levels of total histone H3 (enrichment for H3 is in Supp. Fig. 7) while the level of enrichment of H2A.Z was expressed as percentage of input. (E) Sonication fragmented chromatin was immunoprecipitated using anti-RNAPII antibody or control antibody and analyzed by qPCR using primer specific for the GAPDH promoter region, the HBV enhancer 1 region or for the 5’ core promoter region. Data represents mean ± standard deviation for at least 3 independent experiments, p value was calculated with an unpaired Student’s *t* test with Welch correction (*, P<0.05; **, P<0.01; ***, P<0.001).

To further investigate the role of H2A.Z in cccDNA transcription, and given that H2A.Z facilitates RNAPII recruitment, we performed ChIP experiments on HepG2-NTCP-sec cells silenced for both H2A.Z.1 and H2A.Z.2, or SRCAP, and infected with HBV, using an anti-RNAPII antibody on sonicated chromatin. Silencing of H2A.Z or SRCAP reduces the level of RNAPII associated with viral DNA, but does not affect its binding to the GAPDH promoter (Fig. 4E). Taken together, these results indicate that H2A.Z recruitment by SRCAP facilitates RNAPII recruitment to cccDNA.

To further support our conclusions, and given that chromatin accessibility is influenced by histone modifications and the incorporation of variants such as H2A.Z which modulates nucleosome stability and facilitates nucleosome sliding and eviction, we performed ATAC-seq to assess chromatin accessibility in HepG2-NTCP-sec cells silenced for H2A.Z.1 or H2A.Z.2 individually or in combination or SRCAP. We processed and aligned the ATAC-seq reads to both the human and HBV genomes and found overall that the majority of ATAC-seq reads mapped mostly to introns, enhancers and TSSs as expected (Supp. Fig. 9A). Moreover, ATAC-seq analysis of cellular chromatin confirms that H2A.Z silencing decreases DNA accessibility at known H2A.Z-regulated genes (Supp. Fig. 9B and C). Analysis of cccDNA chromatin accessibility in HBV-infected cells, consistent with previously published results, reveal an overall open chromatin state across the HBV genome, particularly at regulatory regions spanning the enhancer I/X promoter, enhancer II, the basal core promoter, and the PreS/S promoter region (Fig. 5). Alignment with previously published MNase-seq and ChIP-seq data from HBV-infected HepG2-NTCP cells shows that these regions are largely nucleosome-depleted and overlap with RNAPII-enriched areas, and that nucleosome boundaries coincide with increased enrichment of active chromatin marks such as H3K4me3. Silencing of H2A.Z or SRCAP results in a reduction of cccDNA accessibility.

**Figure 5:**
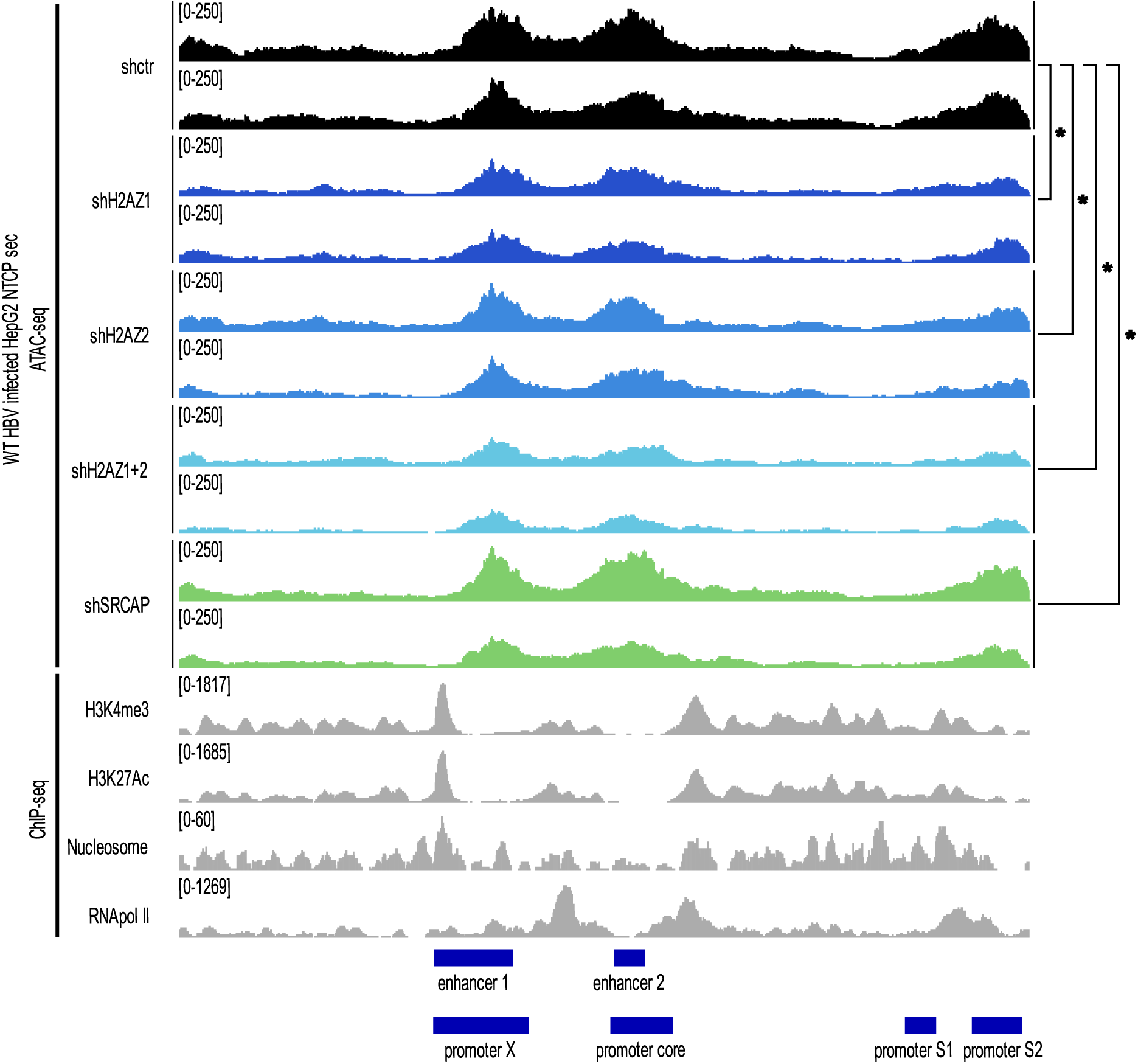
Knocking down H2AZ and SRCAP leads to a decrease in the accessibility of viral DNA. HepG2 NTCP sec cells were transduced with shRNAs targeting H2AZ.1 or H2AZ.2 alone or in combination or targeting SRCAP for 48 hours before being infected with WT HBV at an MOI of 200 Geq/cells. The DNA was tagmented using Tn5 and ATAC-seq libraries were generated and subsequently sequenced using NovaSeq X platform from Illumina. The ATAC-seq signal as well as the ChIP-seq data from Tropberger et al. (H3K4me3, H3K27ac, nucleosome, and polII) were aligned to the linearized HBV genome (genotype D, clone ayw). HBV enhancers and promoters are shown under the mapped sequenced. A one-sided Student’s *t* test was performed to access the significance of the differences in DNA accessibility measured by the ATAC-seq in the different shRNA conditions (*, P<0.01).

Altogether, our results suggest that H2A.Z deposition on cccDNA promotes an open chromatin conformation that facilitates RNAPII recruitment and the establishment of an active transcriptional environment.

### H2A.Z associated factor BRD2 is recruited to cccDNA and regulates HBV transcription

H2A.Z has been shown to recruit Brd2, a member of the BET family of transcriptional co-activators, to active promoters. Mass spectrometry analysis of H2A.Z pulldowns in multiple cell lines identified and revealed a preferential interaction of Brd2 over Brd3 or Brd4 with H2A.Z. This interaction is mediated at least partially by the bromodomain of Brd2^40^. We thus first search whether H2A.Z recruits Brd2 to HBV cccDNA and we performed ChIP analysis using an anti-Brd2 antibody and chromatin from HepG2-NTCPsec cells transduced with lentiviruses coding for shRNAs targeting H2A.Z.1 and H2A.Z.2 or SRCAP and infected by HBV. We found that Brd2 is loaded on the cccDNA and its recruitment to cccDNA decreases in cells silenced for H2A.Z.1 and H2A.Z.2, or SRCAP (Fig. 6A and B).

**Figure 6:**
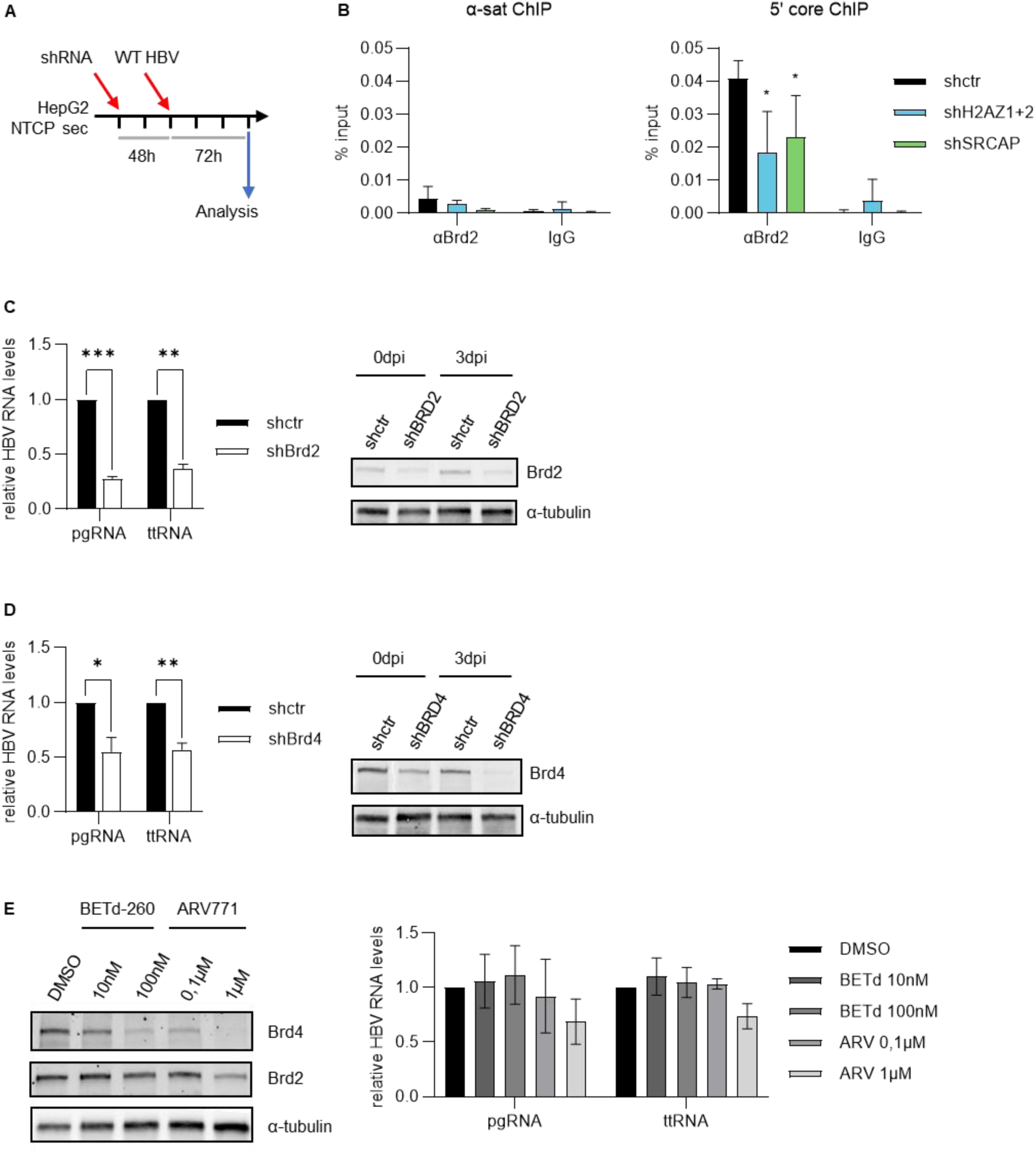
Knocking down Brd2 or Brd4 leads to a decrease in the amounts of HBV RNAs. (A) **experimental design.** HepG2 NTCP sec cells were transduced with shRNAs targeting both H2AZ.1 and H2AZ.2, SRCAP, Brd2 or Brd4 for 48 hours before being infected with WT HBV at an MOI of 200 geq/cell (B) or 100 geq/cell (C) and (D). Chromatin, RNA and proteins were extracted 72 hours post infection. (B) Sonication fragmented chromatin was immunoprecipitated with an antibody targeting Brd2 or normal rabbit IgG and its enrichment for the α-satellite region and the 5’ of the core transcript in the HBV genome was quantified in the immunoprecipitated samples. Data represents mean ± standard deviation for at least 3 independent experiments, p value was calculated with an unpaired Student’s *t* test (*, P<0.05). (C) and (D) Amounts of HBV pgRNA and total HBV RNA (ttRNA) in cell silenced for BRD2 (B) or BRD4 (C) were evaluated by RTqPCR and normalized by the level of GAPDH mRNA. Expression level in cells transduced with shRNA control was set to 1. Data represents mean ± standard deviation for at least 3 independent experiments, p value was calculated with unpaired Student’s *t* test with Welch correction (*, P<0.05; **, P<0.01; ***, P<0.001). BRD2 (B) or BRD4 (C) protein expression in HepG2 NTCP cells transduced by the indicated lentiviral constructs were analyzed by Western blot. Tubulin was used as loading control. (E) HepG2 NTCP sec cells were infected with WT HBV at an MOI of 100 geq/cell for 48 hours before being treated with BETd-260 (BETd) or ARV 771 (ARV) at indicated concentrations. Total RNA and proteins were extracted 24 hours post treatment. HBV pgRNA and total HBV RNA (ttRNA) were evaluated by RTqPCR and normalized by the level of GAPDH mRNA (right pannel). Expression level in control cells (mock treated with DMSO) was set at 1. Data represents mean ± standard deviation for at least 3 independent experiments. Protein levels of BRD2 and BRD4 were analyzed by Western blot. Tubulin was used as loading control (left panel).

To next determine whether Brd2 contributes to HBV transcriptional activation, HepG2-NTCP-sec cells were first transduced with lentiviruses expressing an shRNA targeting Brd2 and then infected with HBV. We observed that Brd2 depletion leads to a decrease in HBV RNA expression (Fig. 6C, Supp. Fig. 2E). As a control, we also depleted Brd4, which has previously been shown to regulate HBV transcription (Fig. 6D, Supp. Fig. 2F).

Interestingly, Brd2 has been evaluated as a therapeutic target in cancer, and several small molecules targeting it are available. We used two proteolysis targeting chimaeric compounds (PROTAC) that induce the degradation of Brd2. We first tested the efficiency of the two drugs (BETd-260 and ARV771) using HepG2-NTCP-sec cells. As shown in Fig. 6E, BETd-260 and ARV-771 preferentially induce Brd4 degradation, whereas ARV-771 also promotes Brd2 degradation when used at a concentration of 1mM. We next evaluated the effects of these compounds in HBV-infected HepG2-NTCP-sec cells. Interestingly, while BETd-260 has no effect on HBV transcription, ARV-771 treatment at 1 µM leads to a decrease in pgRNA and total HBV RNA levels.

Altogether, ours results suggest that Brd2 is recruited on the cccDNA by H2A.Z and participates in HBV transcriptional activation representing thus a potential therapeutic target.

## DISCUSSION

Chromatin organization plays a central role in regulating essential cellular processes such as transcription, replication, and DNA repair. In the context of HBV infection, the mechanisms governing histone deposition onto viral DNA remain incompletely understood, although nucleosome organization on cccDNA has been characterized in several HBV transcription models, where chromatinization was shown to influence HBV transcription^10,11^. However, despite these insights, the contribution of histone variants to the organization and transcriptional regulation of HBV cccDNA remains largely unexplored. In this study we investigated the role of histone variant H2A.Z in HBV replication. We first showed that acetylated H2A.Z is incorporated into transcriptionally active cccDNA and that H2A.Z.1 and H2A.Z.2 are required for efficient HBV transcription. Using gene silencing, we identified SRCAP as the chaperone responsible for H2A.Z deposition on cccDNA. ATAC-seq and ChIP-qPCR analyses revealed that H2A.Z occupancy correlates with increased chromatin accessibility at HBV promoters and enhancers and with enhanced RNA polymerase II recruitment. Furthermore, we demonstrate that H2A.Z regulates HBV transcription through the recruitment of Brd2, which we identified as a potential therapeutic target using silencing approaches and bromodomain inhibitors. Our data also pointed out to a role of H2A.Z on cccDNA establishment since we observed a decrease in HBV cccDNA level in HepG2 NTCP-sec cells silenced for H2A.Z or SRCAP.

Using a multiplexed ddPCR approach (ddOTS) developed to differentiate and quantify the level of the different HBV RNAs, we showed that all HBV promoters are deregulated upon H2A.Z or SRCAP silencing. The downregulation of all HBV transcripts is correlated with a global decrease of DNA accessibility along the HBV genome including the four promoter/ regulatory regions upon SRCAP or H2A.Z.1 and H2A.Z.2 silencing. Using ChIPqPCR we observed the deposition of H2A.Z on different regions on the cccDNA but its precise positioning at HBV promoters or enhancers or both remains to be determined. Indeed H2A.Z has been shown to regulate both the activity of enhancer and promoter by facilitating chromatin accessibility to RNAPII and transcription factors^32,41^. Deciphering more precisely the composition of nucleosomes decorating the cccDNA could be of interest to better understand HBV transcriptional regulation. Recent work, using silencing approaches, revealed that H3.3 which is loaded by HIRA on the cccDNA and is required for HBV transcription^6,19^. H3.3 and H2A.Z are known to interact with each other and the H3.3/ H2A.Z double variant containing-nucleosome has been shown to mark active promoters and regulatory regions facilitating transcripion^32,42,43^. Moreover it has been shown that SRCAP and HIRA cooperate for the deposition of histone variants H3.3 and H2A.Z on poised genes in mESCs suggesting the existence of a crosstalk between the two chaperones^34^. It will be thus interesting to further determine the composition of nucleosomes on the cccDNA and whether the chaperones HIRA and SRCAP cooperate to activate HBV transcription.

In our study we observed that the depletion of SRCAP leads to the decrease of H2A.Z loading on HBV cccDNA suggesting that SRCAP is involved in HBV transcriptional regulation in part via the deposition of H2A.Z. H2A.Z histones can be deposited by a second major chromatin remodeler complex: the p400/TIP60/Nu4A (EP400) complex. In our study we depleted VPS72 which specifically interacts with H2A.Z and observed, as in the depletion of SRCAP, a reduction in the amount of HBV RNAs. Our results with VPS72 given its function in both SRCAP and EP400 complexes as the subunit involved in the deposition of H2A.Z further support a role for H2A.Z in a transcriptionally active cccDNA. A study from Nihituji and collaborators focusing on TIP60/EP400 complex found that it represses HBV preC/pgRNA transcription^44^. In this study the authors did not explore the impact of TIP60/EP400 on H2A.Z deposition. Interestingly, several studies have questioned the role of TIP60/EP400 in H2A.Z deposition, as EP400 depletion results in only a minor decrease in H2A.Z loading^42,45^. In fact, in the study of Pradhan and collaborators, silencing of EP400 impacts more strongly H3.3 deposition^42^. TIP60/P400 may thus not be involved in H2A.Z deposition onto HBV cccDNA. Alternatively, we can hypothesis that SCRAP and TIP60/EP400 participate in fine tuned regulation of HBV transcription involving H2A.Z. It has been shown that while the proximity of H2A.Z containing nucleosome to the TSS influence gene transcription, the positioning upstream or downstream of the TSS have opposing effects for the regulation of epithelial genes during EMT transition. Thus in this context, H2A.Z is removed from the −1 position and relocated at +1, wich correlates with repressed transcription^46^. Whether the role of TIP60/EP400 on HBV cccDNA transcriptional regulation is mostly limited to the acetylation function of the complex will required furher investigation.

Consistent with its role in nucleosome dynamics and DNA accessibility, knockdown of H2A.Z or its chaperone SRCAP reduces HBV cccDNA accessibility in HepG2-NTCP-sec cells which correlates with a decreased loading of RNA polymerase II onto the cccDNA. These results are in line with previous reports showing that H2A.Z promotes a chromatin environment favorable to the recruitment of transcriptional machinery and chromatin remodelers^24,25,32,47^. Moreover, we found that transcriptionally active cccDNA is enriched in acetylated H2A.Z (H2A.Zac), a modification known to correlate with gene activation^36,37,48,49^. Finally, we also observed that Brd2 is recruited to the cccDNA via H2A.Z. Notably, Brd2 has been identified as an H2A.Z interactor whose recruitment is enhanced by the acetylation of histones H4 and H2A.Z^20,40,49–51^. The recruitment of Brd2 may in turn provide a platform for the assembly of histone acetyltransferase (HAT) complexes, thereby promoting further histone acetylation and facilitating the subsequent recruitment of RNA polymerase II^20,40,49^. cccDNA-bound H2A.Zac may thus contribute to HBV transcription by promoting nucleosome destabilization and establishing an active chromatin state^10,12,17^. Although we did not identify the histone acetyltransferase responsible for H2A.Z acetylation in this study, H2A.Z can be acetylated by several histone acetyltransferases, including p300 and KAT2A, both of which have been reported to positively regulate HBV transcription^16,52^. The identification of Brd2 as a coactivator required for HBV transcription is significant, as it may represent a promising therapeutic target. Brd2 is currently considered a potential target in several cancers, and BET inhibitors are already being evaluated in clinical trials. However, the development of more selective compounds is still needed, since available inhibitors target all members of the BET family (BRD4, BRD2, BRD3, and BRDT).

While we found that silencing of H2A.Z.1 and H2A.Z.2 represses HBV transcription upon HepG2-NTCPsec infection, we observed that only H2A.Z.2 silencing correlates with HBV repression in PHH. The role of H2A.Z for HBV transcription in PHH is further supported by the silencing of SRCAP that represses HBV transcription. However it remains unclear why only H2A.Z.2 is required for HBV transcription in PHH. It is possible that in normal liver and in PHH the two isoforms have different functions. Indeed, while the two isoforms differ only by 3 aa and share overlapping functions, they can also regulate specific set of genes or even have antagonist functions^29,31,40,50,53,54^. Recently Tang and collaborators analyzed 116 paired HCC and non tumor adjacent tissues and found that H2A.Z.1 and H2A.Z.2 are overexpressed in tumor tissues. While both isoforms regulate common genes involved in cellular proliferation, H2A.Z.1 regulates specifically genes involved in midzone and microtubule function while H2A.Z.2 controls the expression of genes involved in RNA splicing and export^55^. The reason of this differential activity is unknown but could be due to a difference of structure between H2A.Z.1 and H2A.Z.2, a difference in abundance or localization on the chromatin (i.e. promoter, enhancer or gene body) or a difference in the recruitment of effector proteins^31,56,57^. Liver zonation, where depending on the position of the hepatocytes in the liver different set of genes is expressed, could contribute for the differential activity of H2A.Z^58^. We can speculate thus that depending on the positioning of an hepatocyte population in the liver the respective level of the two H2A.Z isoforms or of their specific interacting partners can differ, leading to a differential function of the two histones variant isoforms.

Interestingly, Belotti and colleagues recently reported that H2A.Z knockout has no detectable effect on transcription in terminally differentiated muscle cells, thereby questioning the role of H2A.Z in postmitotic cells^59^. This observation contrasts with our findings, which indicate that H2A.Z contributes to HBV transcription. Our results are consistent, however, with those of Zhang et al. (2021, Nature Communications), who showed that in the adult mouse liver, transcription is associated with open chromatin, H3K4me3, and H2A.Z loading^60^. Taken together, these observations suggest that the impact of H2A.Z depletion may vary across tissues, highlighting a complex tissue-specific role for H2A.Z in the regulation of gene expression.

In the course of our experiments, we also observed that in hepG2-NTCP-sec infected cells, knockdown of H2A.Z or its chaperone SRCAP correlates with a decrease in HBV cccDNA levels. This observation suggests that H2A.Z might also play a role earlier in the HBV life cycle, during the establishment of cccDNA. Indeed, H2A.Z has been implicated in various DNA repair pathways and in DNA replication (reviewed in Diekmüller et al., 2025)^61^. Interstingly H2A.Z/H3.3 nucleosomes have been reported to facilitate the excision of uracil, implicating these double-variant nucleosomes in DNA repair^62^. Recently Locatelli and colleagues demonstrated that the histone variant H3.3 is loaded onto viral DNA by HIRA very early after infection and is required for cccDNA establishment^6^. In our study, we did not assess the kinetics of H2A.Z loading on the cccDNA and we do not know whether it occurs concomitantly with cccDNA formation. It will be interesting to determine whether both histone variants cooperate for cccDNA formation. Of note, we cannot exclude the possibility that H2A.Z silencing, by altering cccDNA chromatin organization, may also impair cccDNA stability. Future studies will be essential to distinguish between these two hypothesis.

This work therefore broadens our understanding of how HBV co-opts host chromatin mechanisms to establish and maintain transcription. It identifies H2A.Z and the SRCAP complex as key determinants of HBV transcriptional activity and highlights Brd2 as a novel, potentially druggable activator of HBV transcription.

## DATA AVAILABILITY

Previously published datasets used in this study are the following: HepG2 ChIP-seq data from Encode project – H2A.Z (ENCFF757BHF), H3K27ac (ENCFF370SKW), H3K4me3 (ENCFF909WTX), H3K4me1 (ENCFF915OIW), RNA Pol II S5p (ENCFF827WBJ); mononucleosome and ChIP seq data from HepG2-NTCP cells infected with HBV was from Tropberger et al (GSE68402); RNA-seq data from HepG2 cells treated with H2A.Z shRNA was from Yuan et al SRA (PRJNA668278); ATAC-seq datasets were generated in this study.

## ACKNOWLEDGEMENTS

We thank M. Benkirane for helpful discussions and critical reading of the manuscript. We thank all the members of the Molecular Virology Laboratory for their constructive comments. This work was supported by the MSD Avenir program and the Agence Nationale de la Recherche sur le SIDA, les hépatites virales et les Maladies infectieuses émergentes (ANRS MIE). B. J. was supported by a fellowship from the French Ministry of Research and Technology and from ANRS MIE, N. S. was supported by ANRS and the MSD Avenir program. L. G. was supported by a grant from ANRS MIE. O. L., G. P. and J.D.D. was supported by the MSD Avenir program.

## MATERIAL AND METHODS

### Cells

HepAD38 cells derive from HepG2 cells and contains the HBV genome (subtype ayw) under the control of a tetracycline inducible promoter^63^. HepAD38 cells were maintained in Dulbecco modified Eagle medium/F-12 with 10% fetal calf serum (FCS), 3,5 10–7 M hydrocortisone hemisuccinate, and insulin at 5g per ml. Primary human hepatocytes (PHHs) were purchased from Corning (reference 454541) or were isolated as described previously (Pichard et al., 2006) from donor organs unsuitable for transplantation or from liver resections performed in adult patients for medical reasons unrelated to our research program. Liver samples were obtained from the Biological Resource Center of Montpellier University Hospital (CRB-CHUM; http://www.chu-montpellier.fr; Biobank ID: BB-0033-00031) and this study benefitted from the expertise of Dr Benjamin Rivière (hepatogastroenterology sample collection) and Dr Edouard Tuaillon (CRB-CHUM manager). The procedure was approved by the French Ethics Committee and written or oral consent was obtained from the patients or their families. PHH were were maintained in PHH medium (Corning, reference 355056, hepatocyte culture media kit, 500 ml) according to the manufacturer recommendations. HepG2-NTCP sec+ (sec) cells derive from HepG2 NTCP cells that express the human sodium taurocholate cotransporting polypeptide (NTCP) and were selected for their ability to produce high titters of infectious HBV when infected^64^ sec cells are grown in DMEM containing Glutamax with 10% fetal calf serum (FCS).

### Virus production and infection

For virus production, HepAD38 cells were grown in Williams E medium supplemented with 5% FCS, 7×10^-5^ M hydrocortisone hemisuccinate, 5 mg/ml insulin, and 2% dimethylsulfoxide. HBV particles were concentrated from the clarified supernatant by ultracentrifugation for 3h at 32000g on a 20% sucrose cushion. Titers of enveloped DNA-containing viral particles were determined by immunoprecipitation with an anti-preS1 antibody (gift of C. Sureau), followed by DNA extraction and qPCR quantification of viral RC-DNA using rcDNA primers (Table 3).

For infection, only those enveloped particles containing DNA were taken into account. Sec cells were infected in media containing the relevant MOI of virus, 3% DMSO, and 4% PEG (polyethylene glycol 8000) by spinoculation at 37°C (2000rpm, 1h) when possible. PHH were infected in media containing 100 genome equivalent units (geu) per cell 2% DMSO and 4% PEG. After spinoculation cells were incubated over-night with infection media. For both cell types the infection media was removed and cells were washed with PBS multiple times the next day and until the end of the experiments the cells were maintained in culture with respective media supplemented with DMSO.

### cccDNA purification for Mass spectrometry analysis

8 days post-infection, nuclei were prepared from infected PHH using Nuclear and cytoplasmic extraction kit (G. Bioscience) with minor modifications. Briefly, PHH were washed and scraped into ice cold PBS (containing 5mM of sodium butyrate and protease Inhibitor coktails-Roche). After centrifugation at 1500 rpm for 5 minutes at +4°C, cells were resuspended in recommended buffer (SubCell Buffer-I) and homogenized by 5 to 10 strokes in a Dounce homogenizer and incubated on ice for 10 minutes. Cells were then lyzed by adding a recommended volume of lysis buffer (SubCell Lysis Reagent) and by repeating twice a cycle of 5 second of vortexing and 1 minute incubation on ice. For nuclear extraction, the nuclei were isolated by centrifugation at maximum speed (16 000 g) for 5 minutes. The nuclear pellet was resuspended in cold nuclear extraction buffer, kept on ice and vortexed for 15 seconds every 10 minutes for a total of 60 minutes. The sample was then centrifuged at +4°C for 10 minutes at 16 0000g.

For cccDNA purification, the nuclear extract was loaded onto a preformed discontinuous iodixanol gradient (10-40% in buffer 3). After centrifugation at 37 000 rpm for 3h at +4°C in the SW41 rotor (Beckman Instrument), fractions of 0,5 ml were collected from the top of the tube. HBV DNA from each fraction was detected by q-PCR on 5 µl of each fraction or by a specific dot blot hybridization assay. For dot blot analysis, 50µl of fraction were spotted onto a nitrocellulose membrane then denatured with (NaOH 0.2M; NaCl 1M), neutralized with (Tris-HCl 0.5M, pH 7.4; NaCl 1M), followed by a wash with 2x SSC and fixed by baking for 2 hours at 80°C. HBV DNA was detected with a full-length HBV genomic DNA probe labeled with [α-32P]CTP as described previously [Lambert, 1990 #619]. Quantitative analyses were carried out though of PhosphorImager SI system. Positive fractions are pooled, dialyzed then concentrated using Amicon devices. Proteins were then purified and sent to mass spectrometry analysis.

### shRNA

pLKO vectors containing the listed shRNA sequences (Table 1) were purchased from Sigma Aldrich (MISSION® Lentiviral shRNA) as bacterial stock. After amplification and purification of the plasmids HEK293T cells were transfected with the relevant plasmid as well as one encoding for VSVG (pMD2.G) and one encoding the gal-pol (pSPAX2) as previously described [7]. 3 days after transfection the supernatant was harvested, clarified, and concentrated 100-fold by ultracentrifugation (27000g, 1h30min) through a sucrose cushion (20% m/V). The concentration of VLPs was evaluated ELISA assay against p24 and the cells were transduced with an MOI of 10 viral particles per cell in media containing 4µg/mL of polybrene before being spinoculated as for the infection.

**Table 1:**
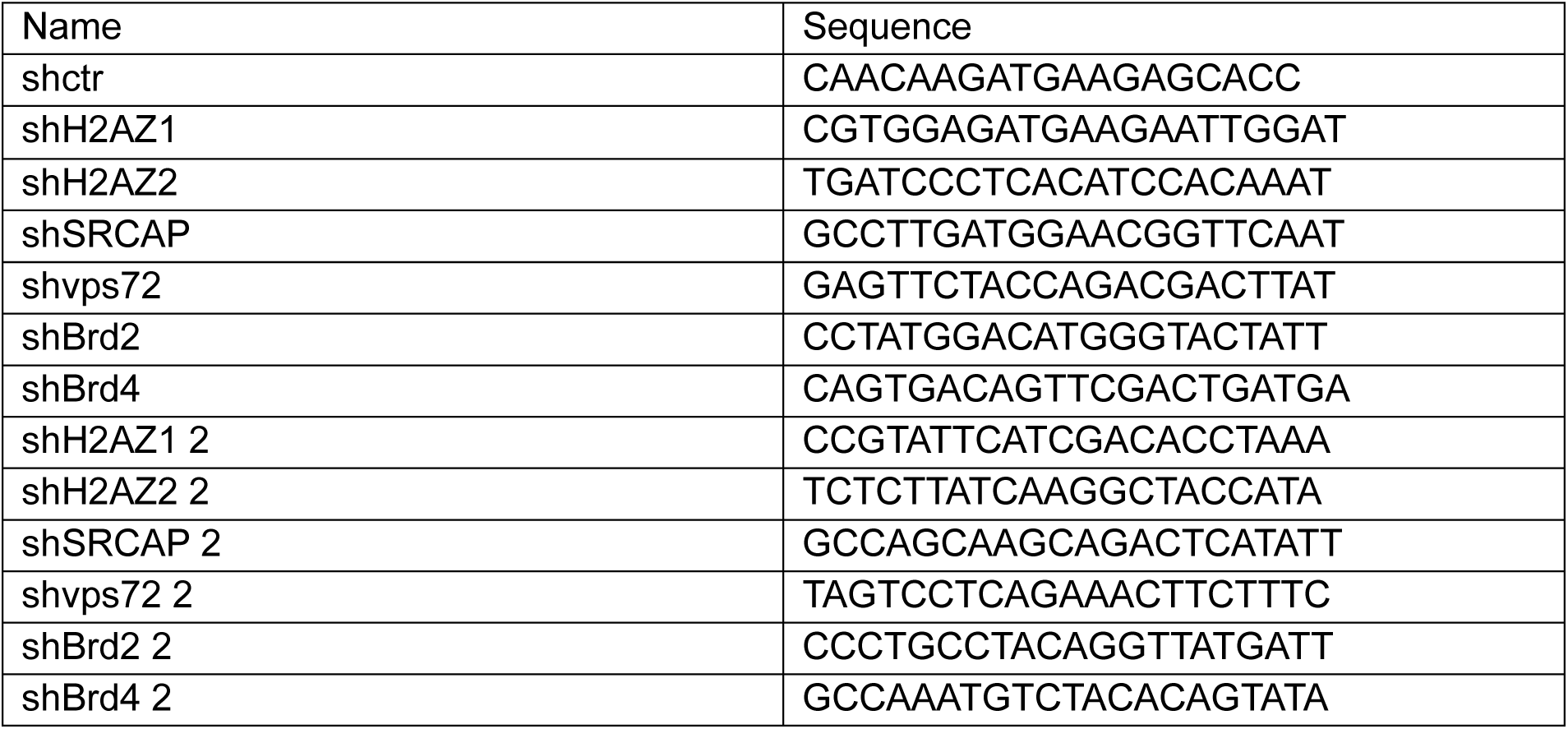
shRNA used in this study.

### Chromatin immuno-precipitation (ChIP)

At the designated time infected HepG2-NTCP sec cells were crosslinked with 1% formaldehyde (room temperature, with rocking) for 10 minutes before being quenched with 0,125M glycine (5min, room temperature, with rocking) and washed twice with cold PBS. The cells were subsequently scrapped in PBS+PIC (Protease Inhibitor Cocktail (200X), Cell Signaling) before being aliquoted and frozen or directly processed for chromatin preparation. For MNAse fragmentation of the chromatin the SimpleChIP® Plus Enzymatic Chromatin IP Kit (Cell Signaling) was used, 5×10^6^ cells were used per IP prep. The manufacturer’s instructions were followed except for that the nuclear membrane was disrupted using a Bioruptor Pico (Diagenode) using the following settings: 20s ON/60s OFF, 4 rounds. For preparation of sonicated chromatin 1×10^7^ cells were resuspended and incubated in lysis buffer 1 [50 mM Hepes-KOH (pH 7.5), 140 mM NaCl, 1 mM EDTA, 10% glycerol, 0.5% NP-40, 0.25% Triton X-100, 1 mM phenylmethylsulfonyl fluoride (PMSF), and PIC] for 10 min at 4°C on a rotation wheel. After centrifugation (1700g, 5min, 4°C), the nuclear pellets were ressuspended in lysis buffer 2 [10 mM tris-HCl (pH 8.0), 200 mM NaCl, and 1 mM EDTA, PIC] and incubated for 10 min at 4°C on a rotation wheel. After centrifugation, nuclei were resuspended and washed twice in shearing buffer D3 [10 mM tris-HCl (pH 7.4), 1 mM EDTA, 0.1% SDS, PIC]. Chromatin was fragmented by sonication with a Covaris E220 Evolution focused ultrasonicator for 8 minutes using the following settings: Peak power (140) / duty factor (5.0) / cycles-burst (200). For both MNAse digested and sonicated chromatin an aliquot of 50µL was used to evaluate chromatin concentration and fragmentation efficiency after reverse crosslinking and DNA extraction using the SimpleChIP® kit as per manufacturer’s instruction. To assess chromatin fragmentation 1µg of DNA was run on a 1% agarose gel. The immunoprecipitation was done using 5µg of MNAse fragmented chromatin or 7µg of sonicated chromatin per IP using the SimpleChIP® kit. Used antibodies were as seen in Table 2. Elution was done at 65°C and 1200rpm for 30 minutes, manufacturer’s instructions were otherwise followed. The DNA samples were then quantified by qPCR using the primers shown in Table 3 and normalized to the input by delta Ct (input Ct-sample Ct) and computed as percentage of input.

**Table 2:** Antibodies used in this study.

| Target | Reference | Dilution |
| --- | --- | --- |
| H2AZ, WB | ab4174, abcam | 1/1000 |
| SRCAP, WB | ESL104, Kerafast | 1/1000 |
| Brd2, WB | A302-583A, Bethyl | 1/1000 |
| Brd4, WB | A700-005, Bethyl | 1/1000 |
| Alpha-tubulin, WB | T5168, Sigma-Aldrich | 1/10000 |
| Goat anti-Rabbit IgG (H+L) Secondary Antibody, DyLight™ 800 | SA5-35571, Invitrogen | 1/10000 |
| Goat anti-Mouse IgG (H+L) Secondary Antibody, DyLight™ 800 | SA5-35521, Invitrogen | 1/10000 |
| H2AZ, ChIP | ab4174, abcam | 2,5µg |
| H2AZAc, ChIP | ABE1363, Merck Sigma | 1µg |
| H3 (D2B12), ChIP | 4620S, Cell Signaling | 2,69µg |
| H3K4me3, ChIP | ab8580, abcam | 2µg |
| H3K9me3, ChIP | ab8898, abcam | 2µg |
| Pol II (F-12), ChIP | sc-55492, Santa Cruz Biotech | 3µg |
| IgG, ChIP | 2729P, Cell Signaling | 2µg |

ChIP, chromatin immunoprecipitation; H+L, heavy and light chains of the IgG molecule; IF, immunofluorescence; WB, western blotting.

**Table 3:**
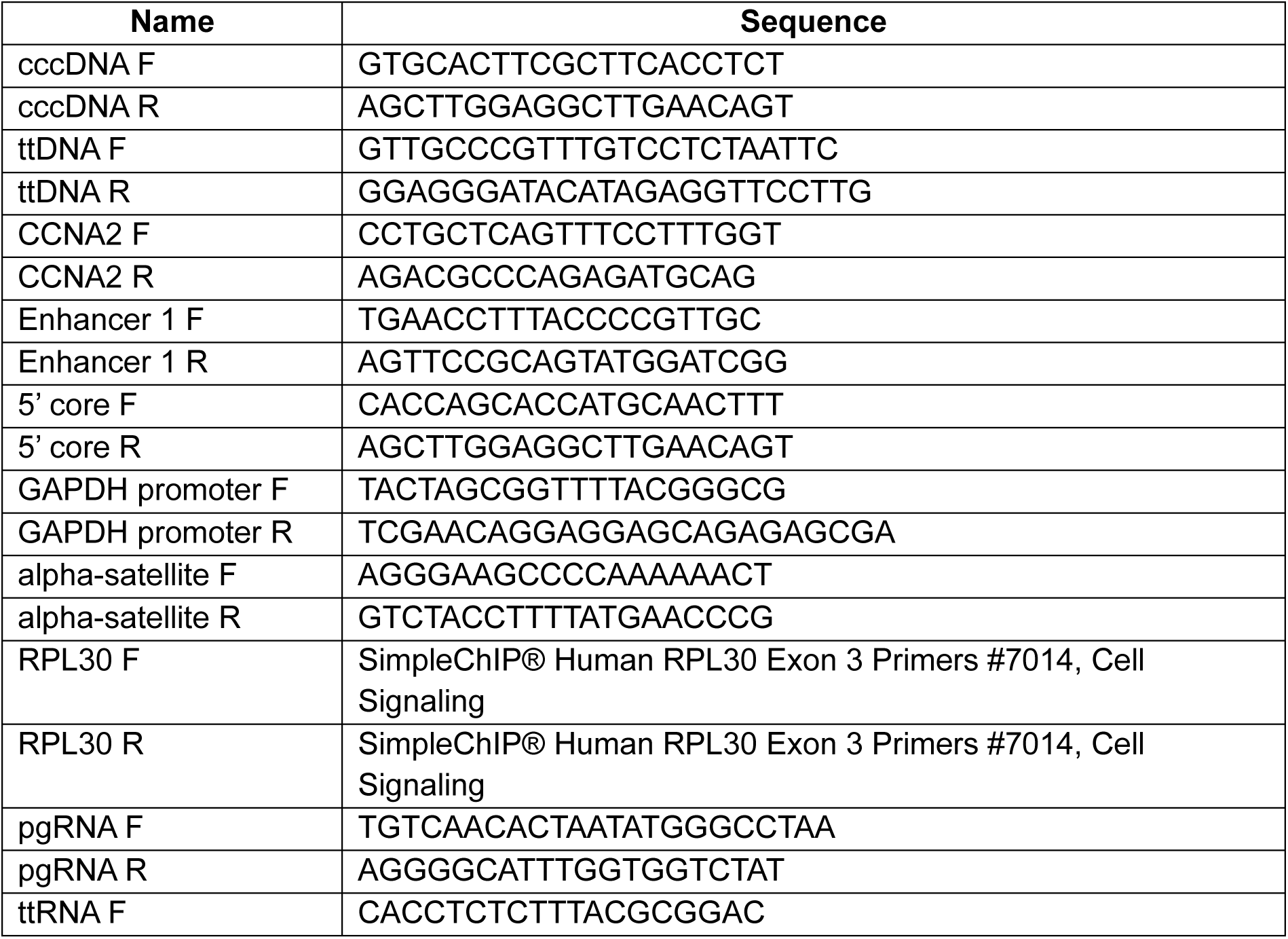
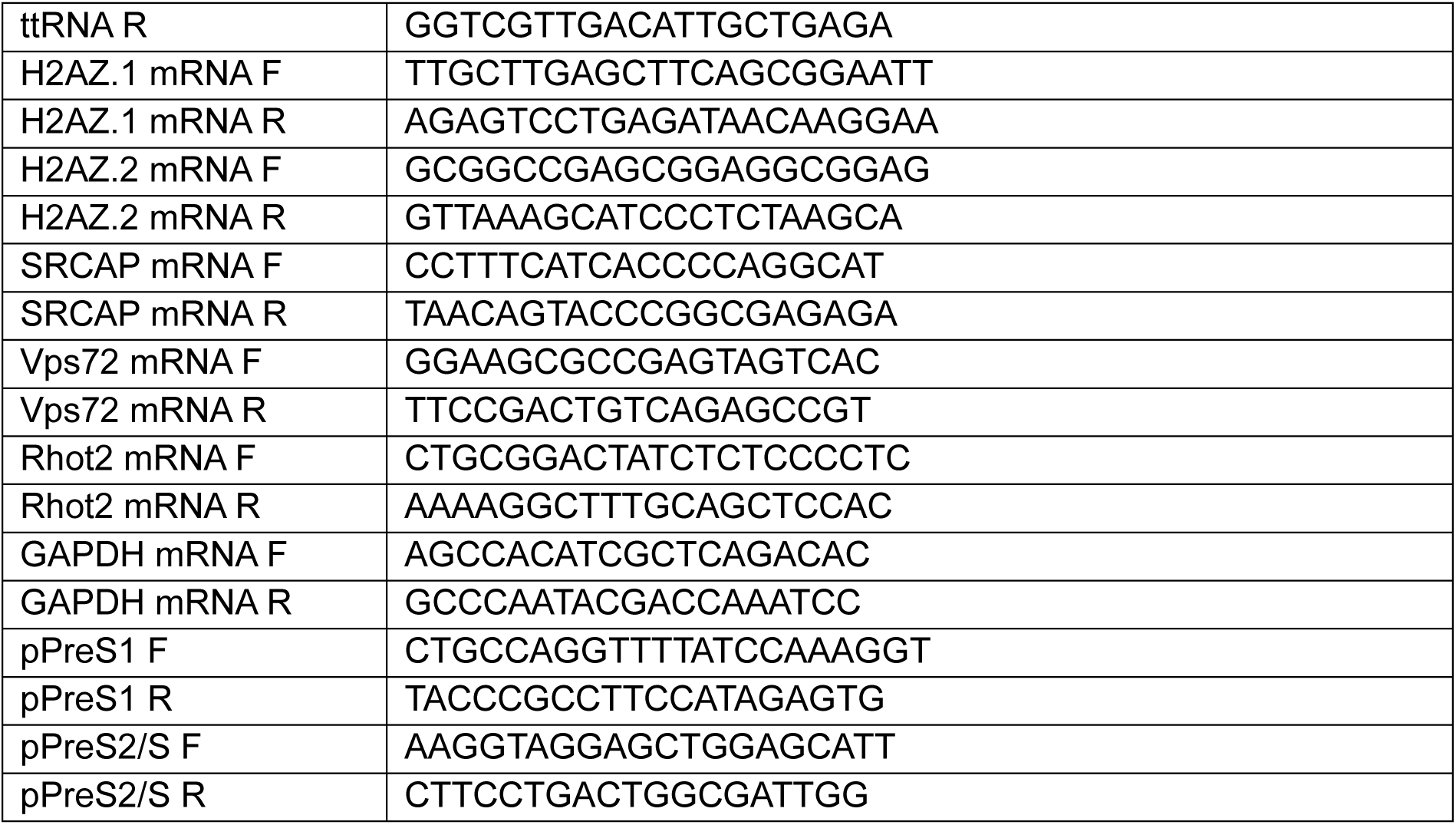
Primers used in this study.

### Quantification of viral nuclear DNA

Cells were lysed in soft lysis buffer (100mM Tris-HCl pH 8.0, 0,375% NP40) and nuclei were pelleted (1min, 14000rpm) to remove the cytoplasmic fraction. The DNA was then extracted using the DNeasy Blood and Tissue kit (QIAGEN) as per manufacturer’s instructions up to the elution where the DNA was eluted in 155µL of AE buffer. To increase the specificity of detection of the cccDNA a fraction of the samples was digested using 20U of Exonuclease I and 25U of Exonuclease III at 37°C for 2h before using cccDNA specific primers, another fraction of the samples was mock digested without the enzymes to quantify total viral DNA and cellular DNA. All the qPCR amplifications were done using the LightCycler® 480 SYBR Green I system (Roche), Table 3.

### Quantitative RT-PCR (RT-qPCR)

Total RNA was prepared using TRIzol reagent (Invitrogen) and TURBO DNase (Ambion). RNA (300ng to 1µg) was retrotranscribed using oligodT primers and RevertAid H Minus M-MuLV reverse transcriptase (ThermoFisher), or SuperScript IV Reverse Transcriptase (ThermoFisher), both according to manufacturer’s instructions. RT-qPCR experiments were carried out as described^54^. For relative quantifications Rhot2 was used as a reference gene because of its low variation coefficient in human liver tumors and cell lines^55^. GAPDH was also used as a reference gene for primary human hepatocytes and conditions where the expression of Rhot2 was affected. HBV RNA and human transcripts were detected using the primer and probes present in Table 3.

### Western Blot

Cells were lysed in RIPA buffer before being sonicated to release chromatin associated proteins (Bioruptor Pico, 30s ON 30s OFF, 2 cycles). The insoluble fraction was then pelleted (16000g, 5min, 4°C) and the supernatant recovered. Proteins were quantified and equal amounts of proteins of each sample were denatured in Laemlli (95°C, 5min) and loaded in Mini-PROTEAN® TGX™ Precast Protein Gels (BioRad). Proteins were then transferred to nitrocellulose membranes using the TransBlot Turbo Transfer System (BioRad). The membrane was then blocked with 5% milk in TBS-T (Tris EDTA ph8.0 10mM, NaCl 150mM, TWEEN 20 0,1%). Membranes were then probed using the indicated primary antibodies and their corresponding secondary antibodies (Table 2) before being imaged using the ChemiDoc Imaging system (BioRad).

### ATACseq

The ATAC-seq method used was based in the Omni-ATAC protocol developed by Corces et al 2017 with minor changes. Briefly, at four days post infection Sec cells were harvested, counted, and 50.000 viable cells were resuspended in 50 ul cold ATAC-Resuspension Buffer (RSB) containing 0.1% NP40, 0.1% Tween-20, and 0.01% Digitonin and incubated 5 min on ice. Lysis was stopped by adding 1 ml of cold ATAC-RSB containing 0.1% Tween-20 to each sample and centrifuging them for 10 min at 4C and 500 RCF. Nuclei were then resuspended in in 50 ul of transposition mixture (25 ul 2x TD buffer, 5 ul transposase (100nM final), 16.5 ul PBS, 0.5 ul 1% digitonin, 0.5 ul 10% Tween-20, 2.5 ul H2O) and tagmented for 45 min at 37C. The DNA was next purified using MinElute PCR Purification Kit (Qiagen) according to manufacturer’s instructions and eluted in 11 ul. For the ATAC-seq libraries preparation 10 ul of the tagmented DNA samples were PCR amplified using NEB Next High Fidelity 2x Mix (NEB) for 5 PCR cycles and then an aliquot of 5ul was used to determine the number of additional cycles by qPCR. After the additional PCR cycles were performed, the libraries were purified using MinElute PCR Purification Kit. To remove excess of adaptors and large DNA fragments a double size selection was performed using Ampure Xp beads (Beckman Coulter) following manufacturer’s guidelines. Libraries were quantified using KAPA library quantification kit (KAPA Biosystems) and fragment size distribution was determined using Bioanalyser (Agilent Technologies). The generated libraries were sequenced using Illumina NovaSeq X Plus paired-end PE150 sequencing. Reads were trimmed using Trimmomatic v0.39 with the parameters ILLUMINACLIP:TruSeq3-SE. fa:2: 30: 10 LEADING:3 TRAILING:3 SLIDINGWINDOW:4: 20 MINLEN:50. Reads were aligned to HBV (HBV.NC_003977.2) and human genome (hg38) using the BWA v0.7.17 aligner. Files were converted into sorted BAM files and filtered using SAMtools (v1.17) Post alignment quality has been made with the R package ATACseqQC (1.30.0). Peaks were called within the aligned reads using MACS2 v2.2.9.1 with the parameters--keep-dup=auto. Statistical significance of differential DNA accessibility was determined using a one-sided paired t-test.

### Statistical analysis

All qPCR results are presented as mean ± standard deviation for at least three independent experiments, p value was calculated with an unpaired t.test with Welch correction. *:P < 0.05, **: P < 0.01, ***: P < 0.001.

## AUTHOR CONTRIBUTIONS

J.D. D. and C. N. designed the study. B.J. and O. L. performed the experiments and analyzed data, with contributions and help from L.G., N. S., G. P., S. G.-C., M. D., and J.D. D. O. H., J.D. D. and C. N. analyzed data and wrote the manuscript with B.J.

## SUPPLEMENTARY FIGURES

**Supplementary Figure 1:**
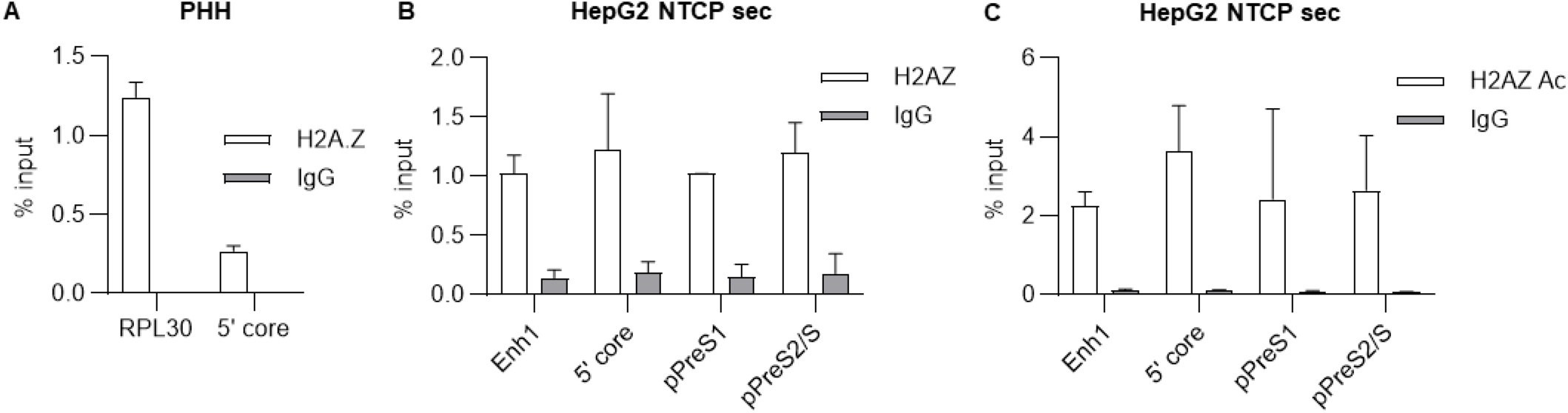
Enrichment of H2A.Z and H2A.ZAc across viral promoters. (A) Primary human hepatocytes were infected at 100 GEQ/cells for 5 days before chromatin extraction. ChIP assays were performed on MNAse fragmented chromatin using anti-H2AZ antibody or nonspecific rabbit IgG as indicated and followed by qPCR analysis with primer specific for the preCore/core promoter region (5’ core) or cellular region (RPL30). Mean ± SD of at least 2 experiments is shown. (B) and (C) HepG2 NTCP sec cells were infected with WT HBV at an MOI of 200 geq/cell for 7 days before chromatin extraction. ChIP assays were performed on MNAse fragmented chromatin using anti-H2AZ antibody, anti-acetylated H2AZ antibody or nonspecific rabbit IgG as indicated and followed by qPCR analysis with primer specific for the enhancer 1 region (Enh1), the preCore/core promoter region (5’ core), the PreS1 promoter region (pPreS1) or the PreS2/S promoter region (pPreS2/S) of the HBV genome. Mean ± SD of at least 2 experiments is shown.

**Supplementary Figure 2:**
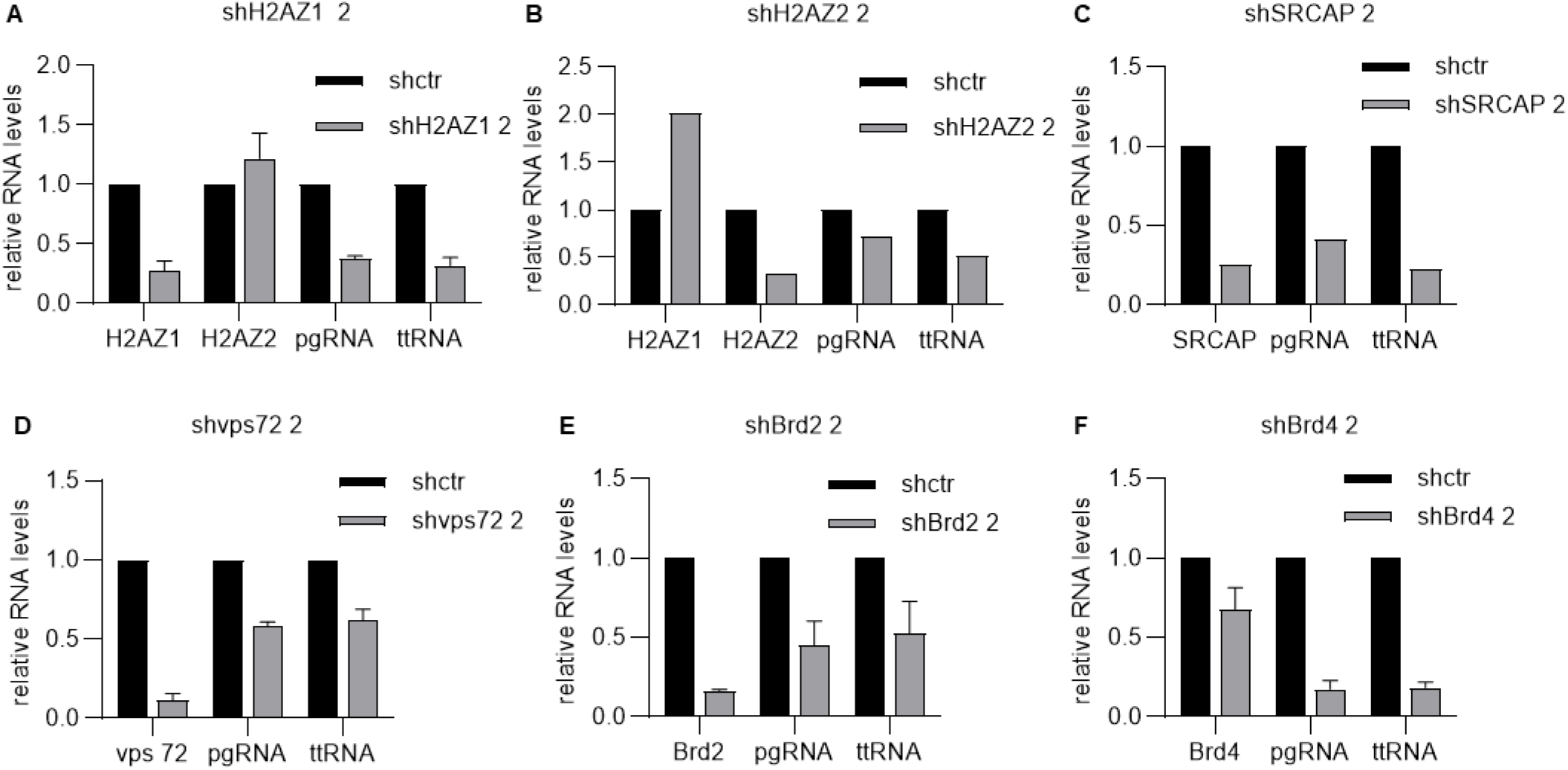
Role of H2AZ1, H2AZ2 and SRCAP on HBV cccDNA transcription. (A) to (F) HepG2 NTCP sec cells were transduced with additional shRNAs targeting H2AZ.1, H2AZ.2, SRCAP, VPS72, BRD2 and BRD4 for 48 hours before being infected with WT HBV at an MOI of 100 geq/cells. Total RNA was extracted at 72 hours post infection. mRNA levels for H2AZ.1 (A), H2AZ.2 (B), SRCAP (C), VPS72 (D), BRD2 (E), and BRD4 (F) as well as HBV pgRNA and total HBV RNA (ttRNA) were evaluated by RTqPCR and normalized by the levels of RHOT2 (A-D) or GAPDH (E-F) mRNA. Expression level in cells transduced with shRNA control was set to 1. Data represents mean ± standard deviation for at least 2 independent experiments or only mean when the experiment was performed once.

**Supplementary Figure 3:**
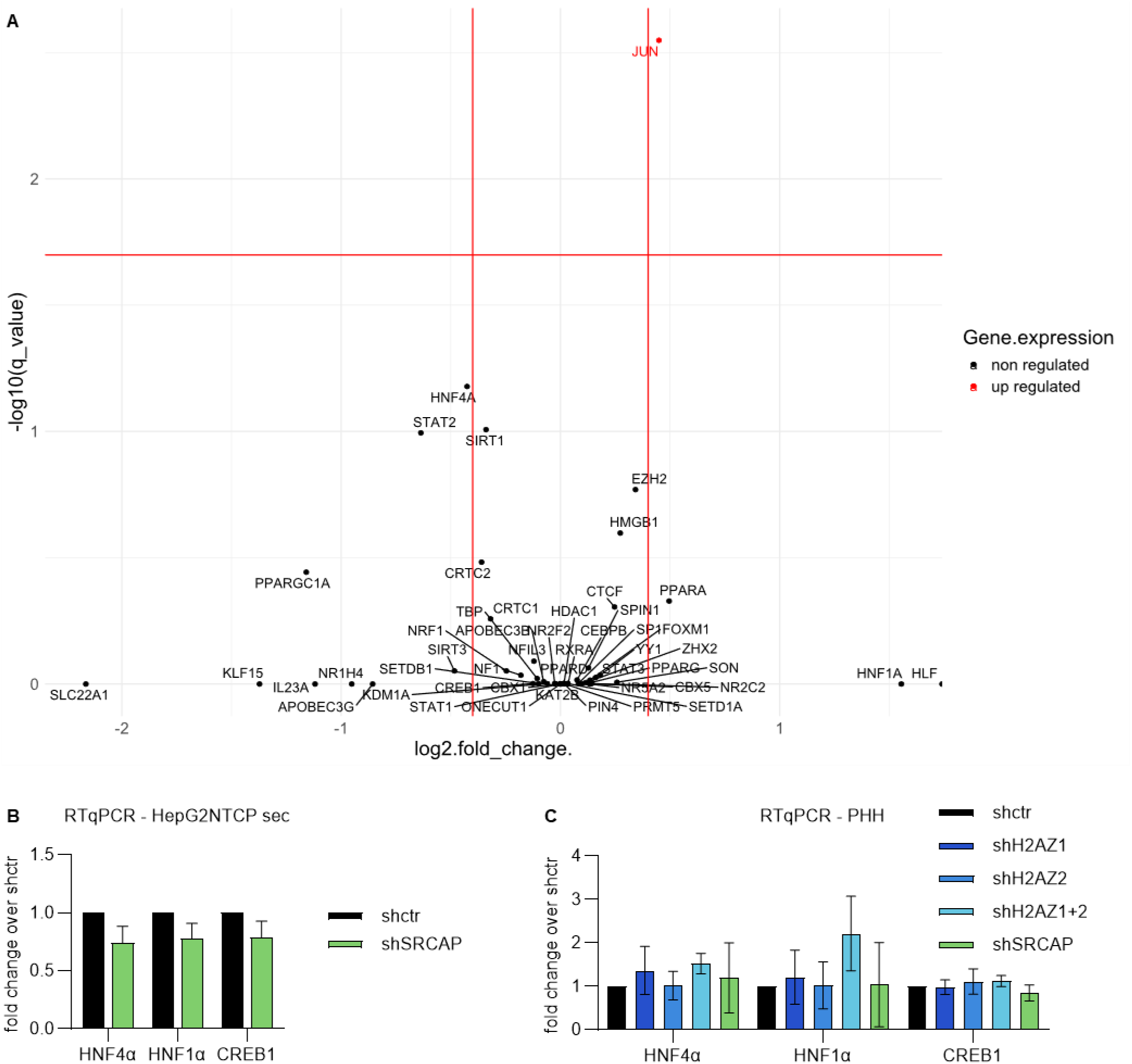
Expression level analysis of known HBV transcriptional regulators. (A) The differential expression of activators and repressors of cccDNA transcriptional activity (Van Damme et al., Front Microbiol, 2021) was calculated based on mRNA-seq data in the presence of shH2AZ from Yuan et al. (Cell Death Dis., 2021). Log2 fold changes in gene expression are plotted against the inverted log10-transformed q values (adjusted p values for multiple testing). The plots shows that the expression of known regulator of hbv transcription are not significantly changed. (B) HepG2 NTCP sec cells were transduced with shRNAs targeting SRCAP or shRNA control (shctr) for 48 hours before being infected with WT HBV at an MOI of 100 geq/cell. Total RNAs were extracted at 72 hours post infection and mRNA levels of the indicated cellular genes were analyzed by RT-qPCR normalized by the level of RHOT2 mRNA. Expression level in cells transduced with shRNA control was set to 1. Data represents mean ± standard deviation for at least 3 independent experiments. (C) Primary human hepatocytes were infected with HBV at MOI of 100 geq/cells for 48 hours before being transduced with shRNAs targeting H2AZ.1 or H2AZ.2 individually or in combination or SRCAP. Total RNA was extracted 5 days post infection and mRNA levels of the indicated cellular genes were analyzed by RT-qPCR normalized by the level of GAPDH mRNA. Expression level in cells transduced with shRNA control was set to 1. Data represents mean ± standard deviation for at least 3 independent experiments.

**Supplementary Figure 4:**
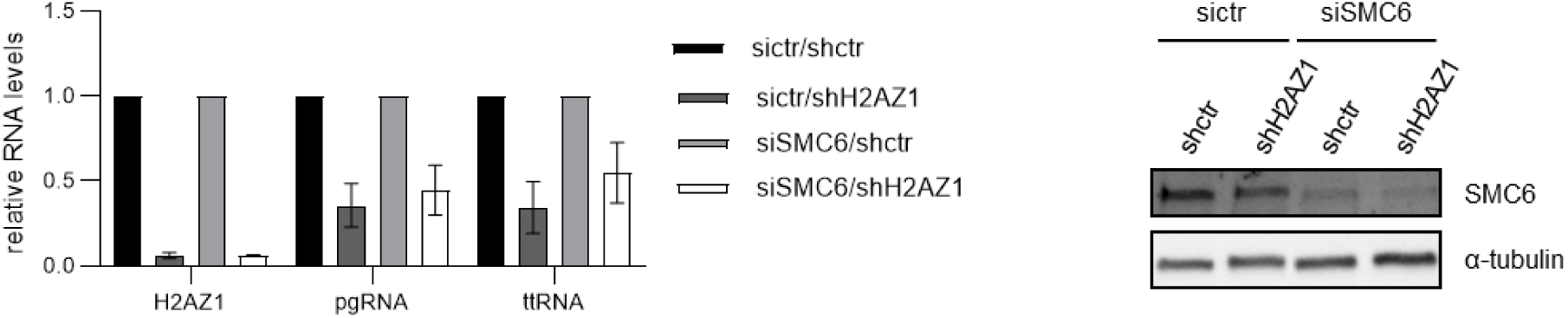
The decrease of HBV RNA levels in cells silenced for H2AZ.1 is not rescued by knocking down SMC5/6. HepG2 NTCP sec cells were transfected with siRNAs targeting SMC6 for 48 hours then transduced with shRNAs targeting H2AZ.1 for 48 hours before being infected with WT HBV at an MOI of 100 geu. RNA and proteins were extracted at 72 hours post infection. Messenger RNA levels of H2AZ.1 as well as pre genomic HBV RNA (pgRNA) and total HBV RNA (ttRNA) were evaluated by RTqPCR, normalized to the levels of RHOT2 mRNA, and the shctr conditions were set to 1. Data represents mean ± standard deviation for 3 independent experiments. Protein level of SMC6 was analyzed by Western blot. Tubulin was used as loading control.

**Supplementary Figure 5:**
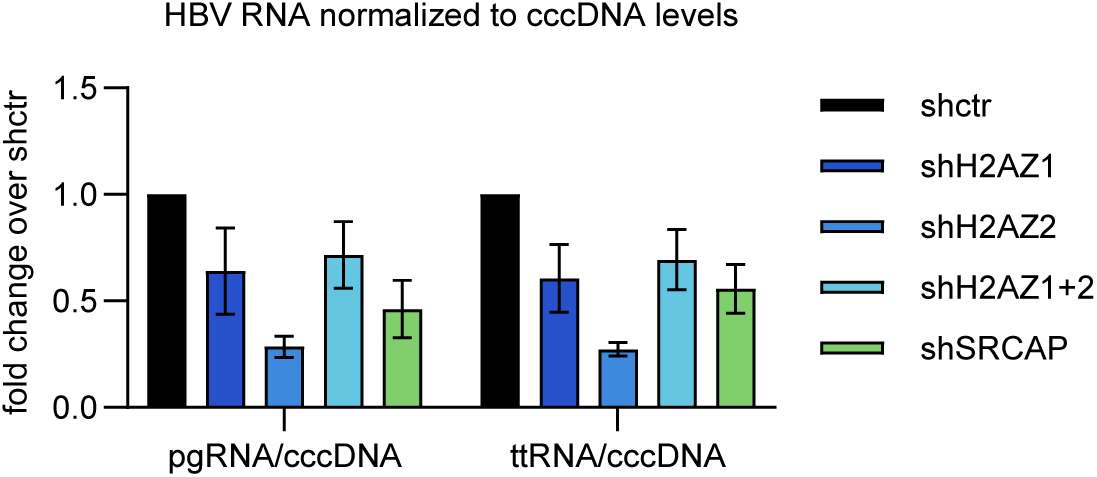
The decrease in HBV cccDNA amounts is not sufficient to explain the decrease in HBV RNA amounts. HepG2 NTCP sec cells were transduced with shRNAs targeting H2AZ.1, H2AZ.2, both or SRCAP for 48 hours before being infected with WT HBV at an MOI of 100 geu. RNA and DNA were extracted at 72 hours post infection. Levels of cccDNA, pre genomic RNA (pgRNA) and total HBV RNA (ttRNA) were measured. The control shRNA treated condition was set to 1 and the levels of RNA were normalized to the levels of nuclear cccDNA. Data represents mean ± standard deviation for 2 independent experiments.

**Supplementary Figure 6:**
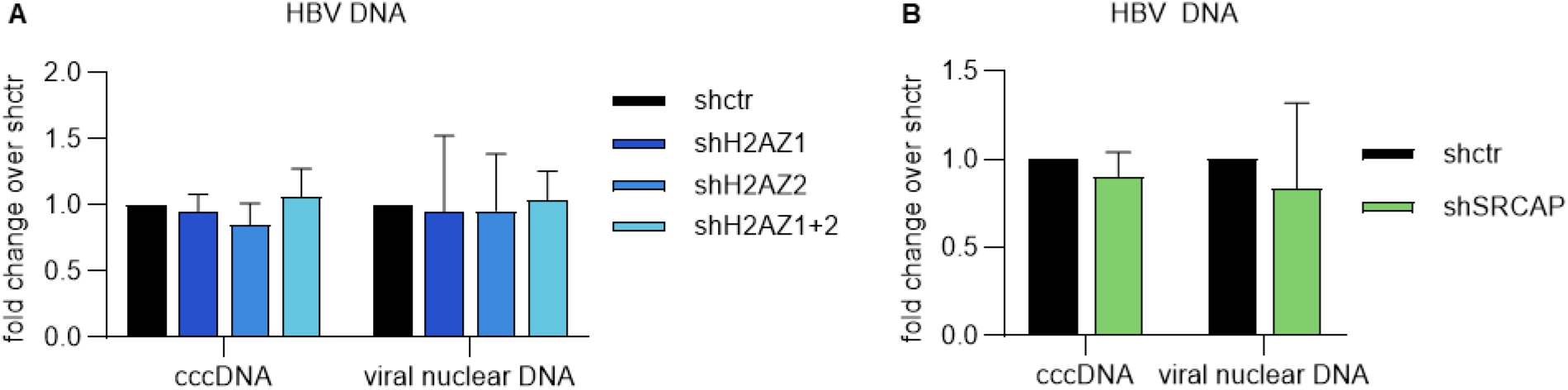
Knocking down H2A.Z and SRCAP after infection of PHH with HBV does not affect the levels of HBV DNA. (A) and (B) Primary human hepatocytes were infected at 100 Geq/cells for 48 hours before being transduced with shRNAs targeting H2AZ.1 or H2AZ.2 or both or targeting SRCAP or shRNA control (shctr). HBV cccDNA and total viral DNA were quantified by qPCR in cells silenced for H2AZ1 or H2AZ2 or both (A) or for SRCAP (B). Viral DNA quantification was normalized to cell number by qPCR amplification of the promoter of cyclin A2. Viral DNA level in cells transduced with shRNA control was set to 1. Data represents mean ± standard deviation for at least 3 independent experiments, p value was calculated with an unpaired Student’s *t* test with Welch correction.

**Supplementary Figure 7:**
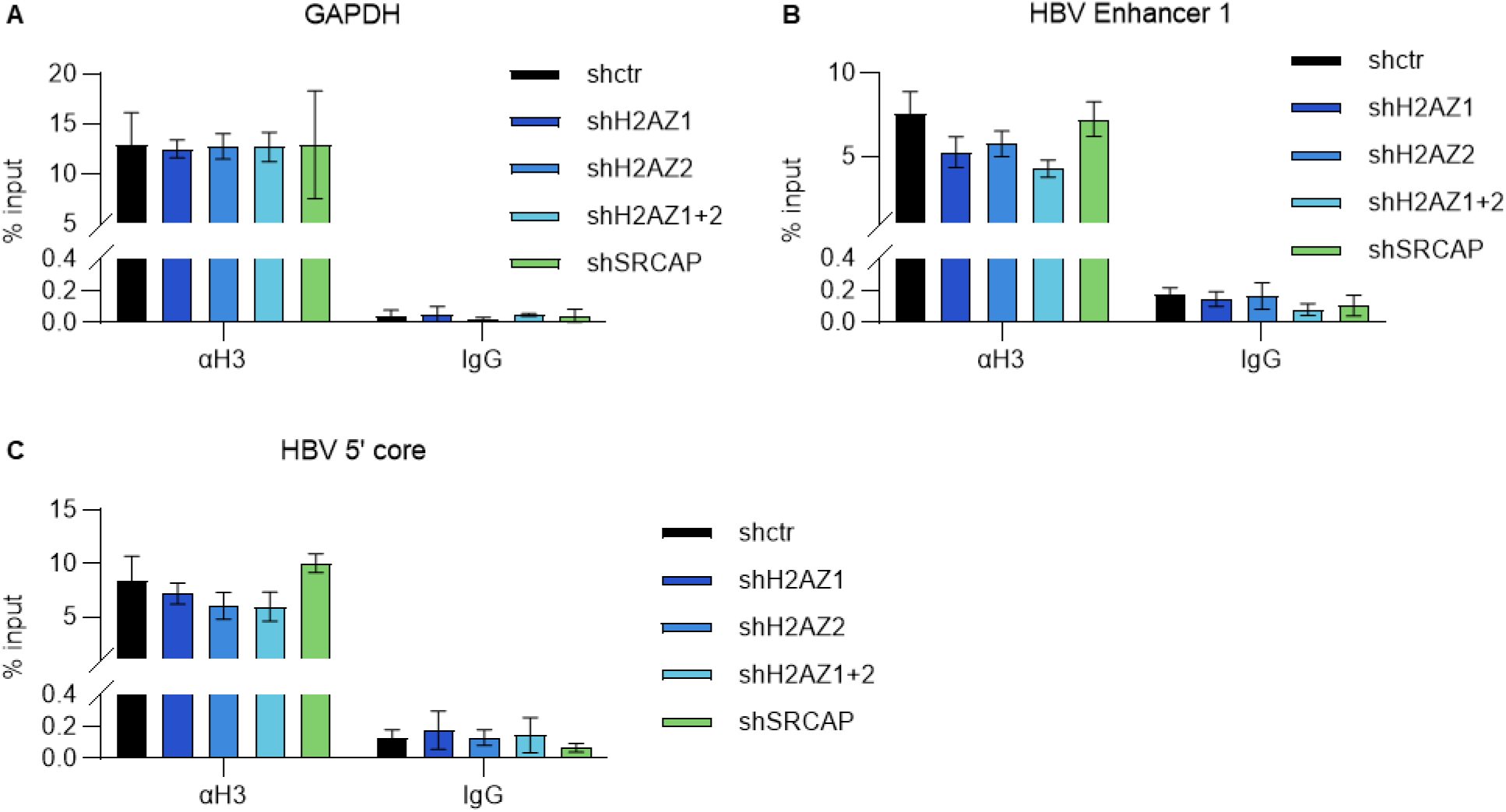
The levels of total histone H3 remain identical in both GAPDH promoter and HBV genome in KD conditions. HepG2 NTCP sec cells were transduced with shRNAs targeting H2AZ.1 or H2AZ.2 or both or targeting SRCAP or shRNA control (shctr) for 48 hours before being infected with WT HBV at an MOI of 200 geq/cell. The cells were crosslinked 72 hours post infection before chromatin extraction. (A) to (C) MNAse fragmented chromatin was immunoprecipitated with antibody targeting H3 or nonspecific IgG and analyzed by qPCR using primers specific for the GAPDH promoter region (A), the HBV enhancer 1 region (B) or for the 5’ core promoter region (C). Data represents mean ± standard deviation for 3 independent experiments.

**Supplementary Figure 8:**
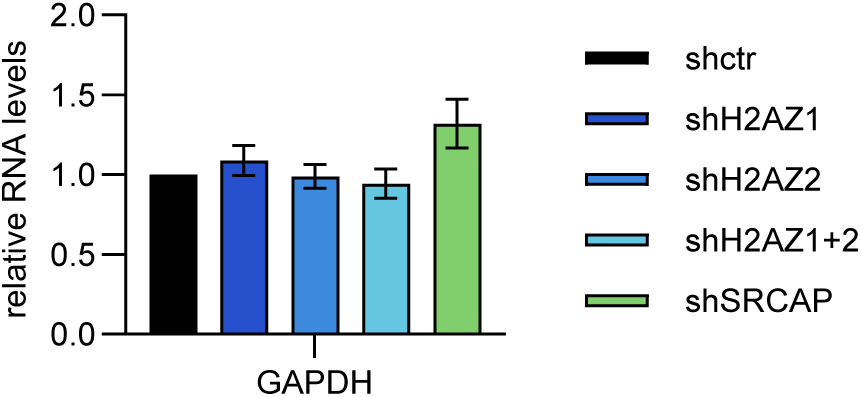
The expression levels of the GAPDH mRNA do not change in the KD conditions. HepG2 NTCP sec cells were transduced with shRNAs targeting H2AZ.1 or H2AZ.2 individually or in combination, or targeting SRCAP or shRNA control (shctr) for 48 hours before being infected with WT HBV at an MOI of 100 geq/cell. GAPDH mRNA level was evaluated by RTqPCR and normalized by the GAPDH mRNA level in the shctr treated condition. Data represents mean ± standard deviation for 3 independent experiments.

**Supplementary Figure 9:**
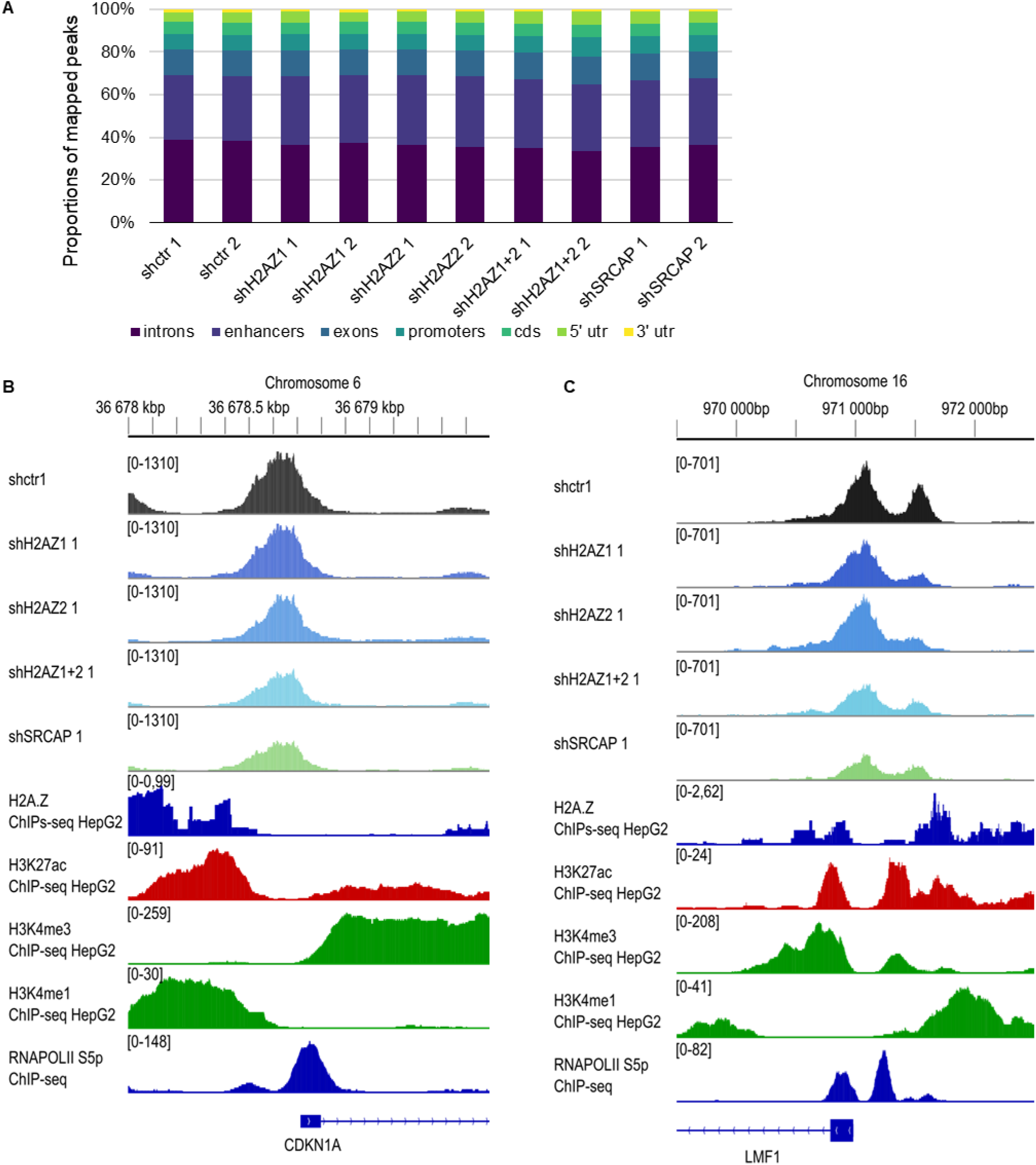
Distribution of ATAC-seq signal relative to different genomic features does not globally change in knockdown conditions while local changes in accessibility can be observed in genes regulated by H2A.Z. (A) Mapped peaks for the human genome were classified according to the genomic features (intron, exon, enhancer, promoter, coding sequence (cds), 5’ utr, and 3’ utr) and their relative abundance in each sample was plotted. Enhancer and promoter positions were taken from the Ensembl database. The others features were retrieved on the annotation made by the Riboraptor package on the Hg38 genome. The peaks positions have been determined using bedtools intersect with an overlap of 100bp with the corresponding feature. One peak can overlap with more than one feature. (B) and (C) The ATAC-seq signal generated in this study and ChIP-seq data from HepG2 cells obtained from the Encode project (H2A.Z, H3K27ac, H3K4me3, H3K4me1 and RNApolII S5P) were aligned to the human genome (GRCh38/hg38). The IGV genome browser profiles of the aforementioned datasets is represented with a window around the TSS of CDKN1A (−1.08 log2 fold change in the shRNA H2A.Z) and LMF1 (−1.07 log2 fold change in the shRNA H2A.Z).

